# Species-specific oscillation periods of human and mouse segmentation clocks are due to cell autonomous differences in biochemical reaction parameters

**DOI:** 10.1101/650648

**Authors:** Mitsuhiro Matsuda, Hanako Hayashi, Jordi Garcia-Ojalvo, Kumiko Yoshioka-Kobayashi, Ryoichiro Kageyama, Yoshihiro Yamanaka, Makoto Ikeya, Junya Toguchida, Cantas Alev, Miki Ebisuya

## Abstract

While the mechanisms of embryonic development are similar between mouse and human, the tempo is in general slower in human. The cause of interspecies differences in developmental time remains a mystery partly due to lack of an appropriate model system^1^. Since murine and human embryos differ in their sizes, geometries, and nutrients, we use *in vitro* differentiation of pluripotent stem cells (PSCs) to compare the same type of cells between the species in similar culture conditions. As an example of well-defined developmental time, we focus on the segmentation clock, oscillatory gene expression that regulates the timing of sequential formation of body segments^2–4^. In this way we recapitulate the murine and human segmentation clocks *in vitro*, showing that the species-specific oscillation periods are derived from cell autonomous differences in the speeds of biochemical reactions. Presomitic mesoderm (PSM)-like cells induced from murine and human PSCs displayed the oscillatory expression of HES7, the core gene of the segmentation clock^5,6^, with oscillation periods of 2-3 hours (mouse PSM) and 5-6 hours (human PSM). Swapping HES7 loci between murine and human genomes did not change the oscillation periods dramatically, denying the possibility that interspecies differences in the sequences of HES7 loci might be the cause of the observed period difference. Instead, we found that the biochemical reactions that determine the oscillation period, such as the degradation of HES7 and delays in its expression, are slower in human PSM compared with those in mouse PSM. With the measured biochemical parameters, our mathematical model successfully accounted for the 2-3-fold period difference between mouse and human. We further demonstrate that the concept of slower biochemical reactions in human cells is generalizable to several other genes, as well as to another cell type. These results collectively indicate that differences in the speeds of biochemical reactions between murine and human cells give rise to the interspecies period difference of the segmentation clock and may contribute to other interspecies differences in developmental time.

## Main

To compare murine and human segmentation clocks *in vitro*, we induced PSM-like cells from mouse embryonic stem cells (ESCs) and human induced pluripotent stem cells (iPSCs) (Fig. 1a), as other groups have recently reported^7–12^. In essence, our PSM induction protocol is based on activation of WNT and FGF signaling as well as inhibition of TGFβ and BMP signaling^9,12^. Prior to the PSM induction, mouse ESCs, which are in the naïve pluripotent state, were pretreated with ACTIVIN A and bFGF and converted to mouse epiblast-like cells (EpiLCs) that possess primed pluripotency as human iPSCs do. To visualize the segmentation clock in the induced PSM, we introduced a HES7 promoter-luciferase reporter^13,14^, detecting clear synchronized oscillations of HES7 expression in both murine and human PSM (Fig. 1b; Supplementary Video 1). Interestingly, the oscillation periods, i.e., the durations for one cycle, were different between the species: mouse PSM oscillated with a period of 122 ± 2 min (mean ± sd) whereas human PSM exhibited a 322 ± 6 min period (Fig. 1c-e). These numbers are consistent with the literature: The period of the murine segmentation clock *in vivo* is 2-3 hours^13,15,16^. While visualizing the segmentation clock in a human embryo is ethically difficult, the human period has been roughly estimated to be 4-6 hours with fixed samples through counting the number of somites, which are periodically formed according to the segmentation clock^17,18^. Thus, we concluded that our induced PSM recapitulates the species-specific periods of the segmentation clock and serves as an ideal *in vitro* platform to investigate the cause of the 2-3-fold period difference between mouse and human.

**Figure 1.**
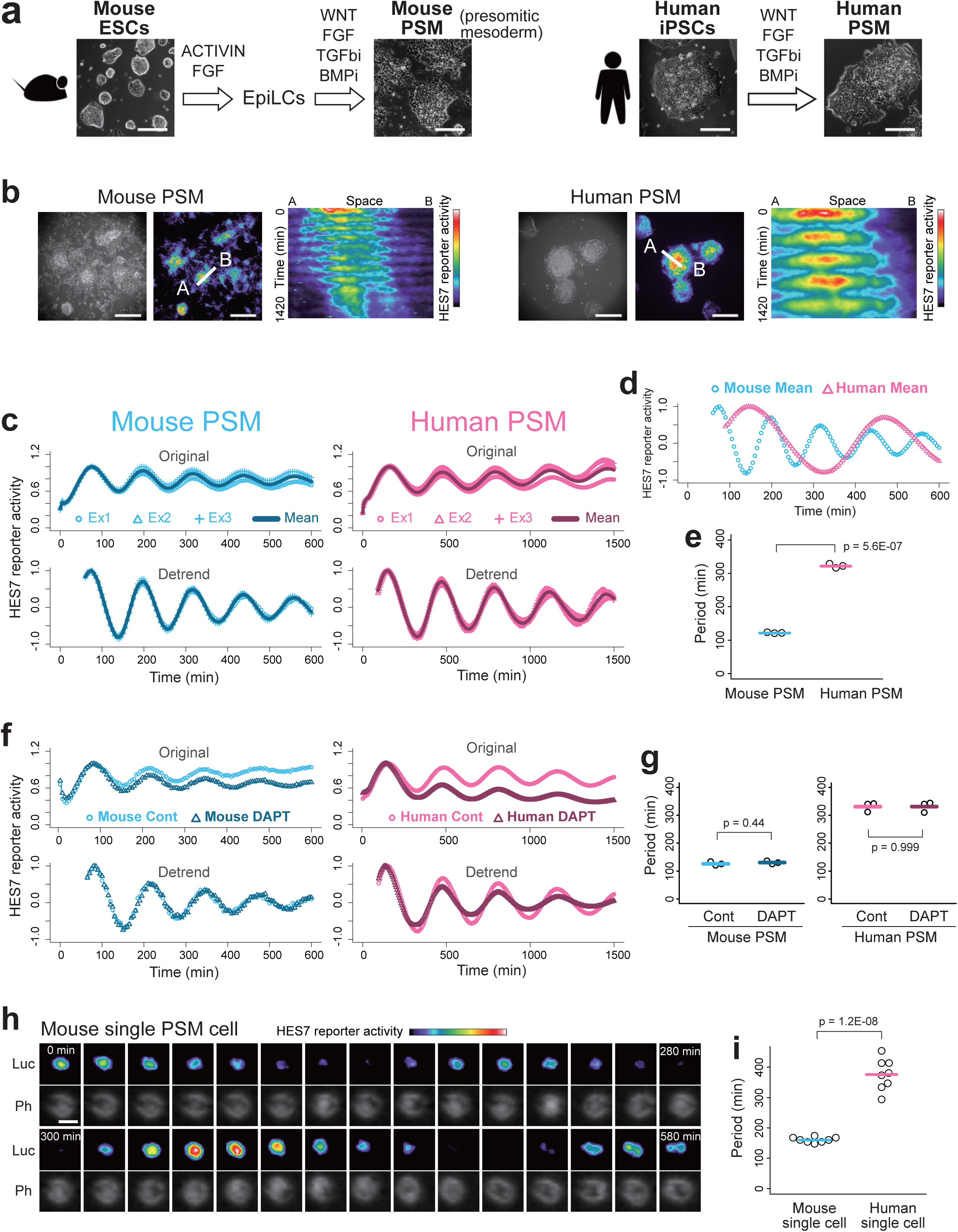
Cell autonomous period difference between murine and human segmentation clocks. **a**, Induction of PSM-like cells. Mouse ESCs were pretreated with ACTIVIN A and bFGF to induce mouse EpiLCs. Mouse EpiLCs and human iPSCs were treated with WNT agonist, bFGF, TGFβ inhibitor, and BMP inhibitor to induce the PSM fate. Scale bars: 200 μm. **b**, HES7 reporter activities in murine and human PSM in time-lapse imaging. Kymographs indicate the spatio-temporal signals along the line between the point A and B. Scale bars: 400 μm. See also Supplementary Video 1. **c**, Oscillatory HES7 reporter activity measured with a luminometer. Original (top) and detrended (bottom) signals of three independent experiments. **d**, Overlay of mean murine and human signals shown in c. **e.** Oscillation periods estimated from c. **f**, Effects of NOTCH signaling on the oscillation periods. Murine or human PSM were treated with NOTCH inhibitor DAPT (10 μM). Representative of three independent experiments. **g**, Periods estimated from f and Supplementary Fig. 1b. **h**, Time-lapse imaging of single cells. Mouse PSM cells were sparsely split, and the oscillatory HES7 reporter activity in a single cell was monitored. See also Supplementary Video 2. Ph: Phase image; Luc; Luciferase image. Scale bar: 20 μm. **i**, Periods of single murine and human PSM cells estimated from Supplementary Fig. 2b, c. All p-values are from two-sided student’s t-test.

The gene regulatory network of the segmentation clock consists of two parts: the intracellular network that gives rise to a cell autonomous oscillation in each cell and the intercellular network that synchronizes the oscillations among neighboring cells (Supplementary Fig. 1a)^19–21^. We therefore first attempted to clarify whether the interspecies period difference stems from the single-cell oscillator or the multicellular synchronized oscillations. Because cell-cell communication through NOTCH-DELTA signaling has been reported to synchronize oscillations among cells by regulating HES7 expression^20,22–24^, we treated both murine and human PSM with a NOTCH inhibitor, DAPT. While the expression level of the HES7 reporter decreased upon administration of DAPT (Fig. 1f, Original), the oscillation period did not change significantly in either species (Fig. 1f, g; Supplementary Fig. 1b). Although WNT and FGF signaling pathways have also been reported to modulate the segmentation clock^25–28^, the existence of high dosages of a WNT agonist, CHIR, and bFGF in the culture medium suggests that cell-cell communication through these pathways should not be crucial for the interspecies period difference. Furthermore, we measured the oscillation period in a sparse cell culture, where cells do not contact each other (Fig. 1h; Supplementary Fig. 2; Supplementary Video 2). Those isolated PSM cells still displayed the 2-3-fold period difference between the species (mouse: 160 ± 9 min; human: 376 ± 51 min) (Fig. 1i), even though the oscillations at the single-cell level were noisier and slower than the population level oscillations (see Supplementary Fig. 2b, c). These results indicate that the period difference of HES7 oscillation between mouse and human is cell autonomous, and that the cause of the interspecies difference should lie in the oscillation mechanism at the single-cell level.

HES7 oscillations have been proposed to arise from a delayed autoregulatory negative feedback loop: HES7, a transcription repressor, directly binds to and inhibits its own promoter with time delays, resulting in an oscillatory expression of HES7 (Supplementary Fig. 1a)^6,14,29,30^. Knocking out other HES family members, such as HES1 and HES5, does not disrupt segmentation in mouse embryos^31^. Since HES7 itself is considered the most critical gene for HES7 oscillation, we first hypothesized that differences in the sequences of HES7 loci between murine and human genomes might lead to the observed oscillation period difference. To test this hypothesis, we swapped HES7 loci between mouse and human (Fig. 2a): the human HES7 locus, which was defined as the sequence including a promoter, exons, introns, and UTRs of HES7, was knocked into the mouse HES7 locus in mouse ESCs (Fig. 2b; Supplementary Fig. 3), and the resulting cells were induced to differentiate into the PSM fate. The homozygous knock-in (i.e., human HES7/human HES7, hereafter referred to as ‘homo swap’) PSM and the heterozygous knock-in (human HES7/mouse HES7, hereafter referred to as ‘hetero swap’) PSM showed slightly longer oscillation periods of 146 ± 7 min and 133 ± 4 min, respectively, as compared with the 124 ± 3 min period of wild-type (mouse HES7/mouse HES7) mouse PSM (Fig. 2c, d; Supplementary Fig. 4a). Considering that the period of wild-type human PSM is 322 min (see Fig. 2d, Wt Human PSM), however, the period extension in homo swap mouse PSM is minor. To confirm this finding, we created knock-in mice containing the human HES7 locus. The homo swap mice appeared largely normal, even though their vertebrae, which are derivatives of somites and therefore subject to the influence of the segmentation clock, displayed minor defects (Fig. 2e; Supplementary Fig. 5). The *ex vivo* measurements of the segmentation clock in homo swap embryos showed ~30 min longer oscillation period as compared with wild-type embryos (Fig. 2f-h; Supplementary Fig. 4b), consistent with the ~20 min period extension in the homo swap samples of induced PSM (see Fig. 2d). However, the 20-30 min period extension in homo swap PSM/embryos is far from the ~200 min period difference between mouse and human, so these results suggest that human HES7 locus in mouse PSM gives rise to an essentially mouse-like oscillation period.

**Figure 2.**
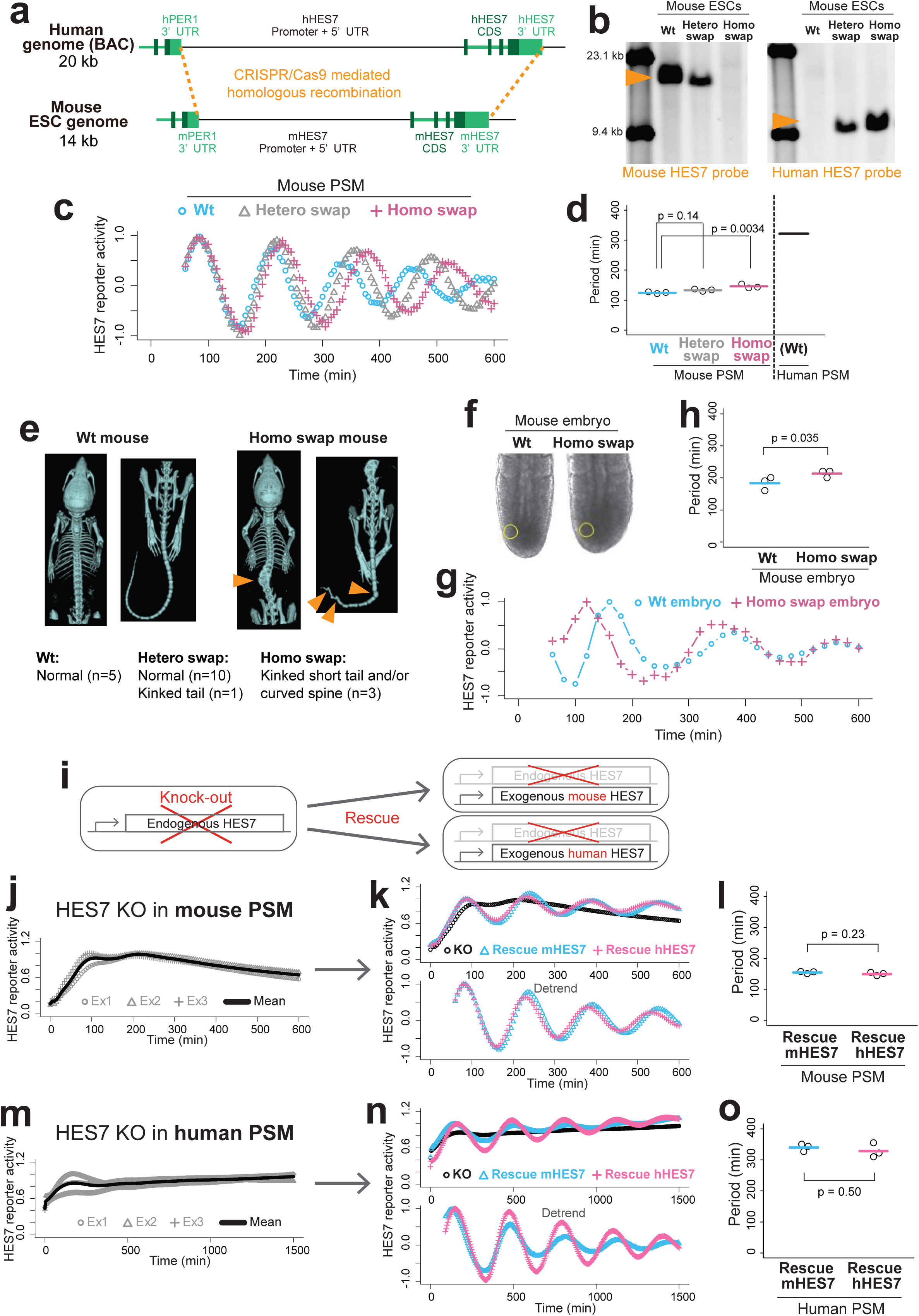
Effects of sequence differences between murine and human HES7 loci on the oscillation period. **a**, Swapping of the human and murine HES7 loci with CRISPR/Cas9-mediated homologous recombination. The HES7 locus was defined as the region ranging from the end of 3’UTR of the adjacent gene, PER1, to the end of 3’UTR of HES7. **b**, Southern blotting of mouse ESCs containing the human HES7 locus with probes against murine (left) and human (right) HES7 sequences. Wt: mouse HES7/mouse HES7; Hetero swap: human HES7/mouse HES7; Homo swap: human HES7/human HES7. **c**, Oscillatory HES7 reporter activity in mouse PSM containing the human HES7 locus. Mean of three independent experiments. **d**, Periods estimated from Supplementary Fig. 4a. The period of wild-type human PSM shown in Fig. 1e is displayed as a control. P-values are from two-sided Dunnett’s test. **e**, Phenotypes of the knock-in mice containing the human HES7 locus. Four-week old mice were scanned with μCT. **f, g**, *Ex vivo* tail bud cultures of the mouse embryos containing the human HES7 locus. The tail buds of E10.5 mouse embryos were cultured (f), and the oscillatory HES7 reporter activity was monitored (g). Signals were averaged within the yellow circle, and a representative of three independent experiments is shown. **h**, Periods estimated from g and Supplementary Fig. 4b. P-value is from two-sided paired t-test. **i-o**, Knock-out (KO) and rescue assay. **j, m**, Endogenous HES7 genes were knocked out in murine (j) and human (m) cells to disrupt HES7 oscillation. **k, n**, The disrupted oscillations were rescued with either an exogenous construct containing a promoter, exons, introns, and UTRs of mouse HES7 (mHES7) or human HES7 (hHES7). Mean of three independent experiments. The mean data of KO shown in j, m are displayed as a control. **l, o**, Periods estimated from Supplementary Fig. 4c. P-values are from two-sided student’s t-test.

One potential defect in our experimental design of interspecies genome swapping is, however, that the swapped HES7 region might not be long enough, and that a crucial sequence for the oscillation period might exist upstream of the HES7 promoter we defined, for instance. To rule out this possibility, we performed ‘knock-out and rescue’ assays (Fig. 2i): The endogenous mouse HES7 gene was first knocked out in mouse ESCs, leading to disruption of the HES7 oscillation in the induced PSM (Fig. 2j). Then the disrupted oscillation was rescued by introducing an exogenous construct containing a promoter, exons, introns, and UTRs of murine or human HES7 locus (Fig. 2k; Supplementary Fig. 4c). Note that these exogenous constructs were integrated into random positions of the genome by transposon vectors, implying that the HES7 regions used for the constructs should be sufficiently long to restore the oscillations. Importantly, both murine and human HES7 constructs restored mouse-like oscillation periods in the mouse PSM (Fig. 2l). We further attempted a ‘complementary’ experiment: we knocked out the endogenous human HES7 gene and rescued the disrupted oscillation with the murine or human HES7 construct in human PSM (Fig. 2m, n). Again, murine and human HES7 constructs were indistinguishable in terms of the restored oscillation period (Fig. 2o). These results collectively indicate that the 2-3-fold period difference between murine and human segmentation clocks is not caused by the sequence differences between murine and human HES7 loci.

We then hypothesized that differences not in the sequences but in the biochemical reaction speeds of HES7 between murine and human cells might lead to the observed oscillation period difference. Since the most important biochemical parameters that affect the oscillation period of HES7 are the degradation rates of HES7 (i.e., δ_m_ and δ_p_ in Fig. 3a) and the delays in the feedback loop of HES7 (τ_Tx_, τ_In_, τ_Tl_, and τ_Rp_ in Fig. 3a)^14,20,29,30,32^, we measured those parameters in both murine and human PSM, exploring which parameter(s) are different between the species. To measure the degradation rate of HES7 protein (δ_p_), we overexpressed either the murine or human HES7 sequence and then halted its expression (Fig. 3b). Interestingly, both murine and human HES7 proteins were degraded more slowly in human PSM as compared with mouse PSM (Fig. 3b, c; Supplementary Fig. 6a), meaning that the changes in the degradation rate depend on the differences not in the HES7 sequences but in the cellular environments (i.e., whether HES7 is hosted in a murine or human cell). The half-life of HES7 protein in mouse was previously reported to be 22 min^29^, consistent with our measurements where half-lives in murine and human PSM were estimated to be 21 ± 0.8 min and 40 ± 4 min, respectively.

**Figure 3.**
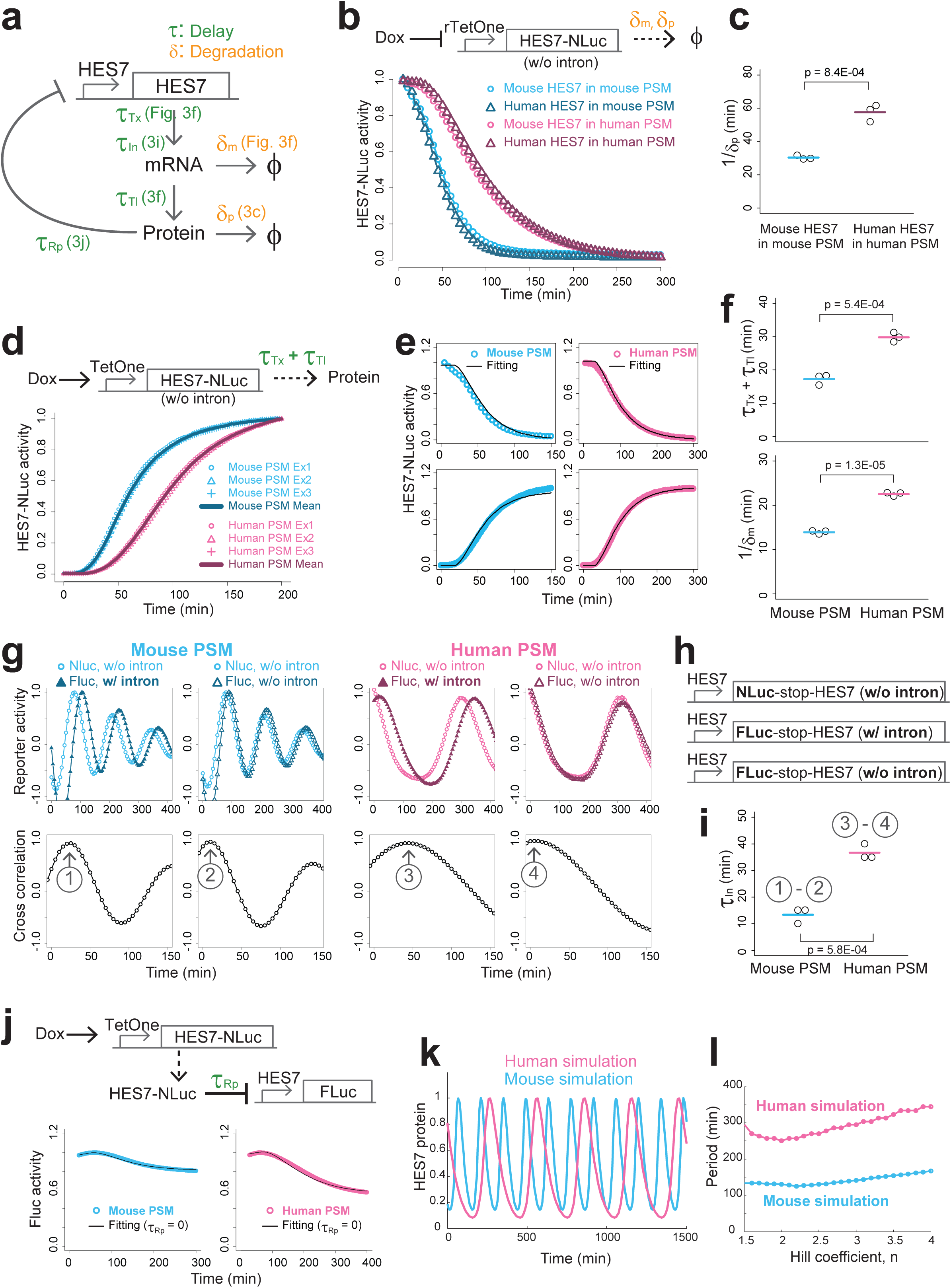
Measuring biochemical parameters of HES7 that determine the oscillation period. **a**, Schematic representation of the negative feedback loop of HES7. The biochemical parameters that determine the oscillation period, i.e., delays and degradation rates, were measured in the indicated panels. τ_Tx_: Transcription delay; τ_In_: Intron delay; τ_Tl_: Translation delay; τ_Rp_: Repression delay; δ_m_: Degradation rate of mRNA; δ_p_: Degradation rate of protein. **b**, Degradation assay of HES7. The transcription of the fusion construct of HES7 and NanoLuc (NLuc) was halted upon Doxycycline (Dox) addition at t = 0 (top), and the decay of NLuc signal was monitored (bottom). Either mouse HES7 or human HES7 sequence was used in murine or human PSM. Mean of three independent experiments. **c**, Inverse degradation rates of HES7 protein estimated from Supplementary Fig. 6a. **d**, Expression delay assay of HES7. The transcription of HES7-NLuc construct was induced upon Dox addition at t = 0 (top), and the onset of NLuc signal was monitored in either murine or human PSM (bottom). **e**, Fitting of the HES7 degradation data shown in b (Mouse HES7 in mouse PSM and Human HES7 in human PSM) (top) and fitting of HES7 expression delay shown in d (Ex1) (bottom). **f**, Transcription/translation delays of HES7 (top) and inverse degradation rates of Hes7 mRNA (bottom) estimated from e and Supplementary Fig. 6b. **g, h**, Intron delay assay of HES7. Three reporter constructs were used (h). Stop: stop codon. Dual reporter assays with NLuc and Firefly luciferase (FLuc) were performed (g, top), and the cross correlation functions of NLuc and FLuc signals were calculated (g, bottom) in either murine or human PSM. Mean of three independent experiments. **i**, Intron delays of HES7 estimated from Supplementary Fig. 7. **j**, Repression delay assay of HES7. The transcription of HES7-NLuc was induced upon Dox addition at t = 0, and the induced HES7 protein repressed the transcription of the FLuc reporter in either murine or human PSM (top). Fitting of the repression data with the parameter repression delay = 0 (bottom). Mean of three independent experiments. **k**, Simulating HES7 oscillations with measured biochemical parameters. Hill coefficient n = 3. The other parameters are summarized in Supplementary Table 1. **l**, Periods estimated by computing the power spectra of simulated oscillations with different values of repression Hill coefficient. All p-values are from two-sided student’s t-test.

To measure the delay caused by the transcription and translation of HES7, we induced the expression of HES7 and estimated the onset time by fitting the results to a standard gene expression model in which transcription and translation are assumed to occur in a linear manner with the corresponding delays (Fig. 3d, e; Supplementary Fig. 6b). The transcription and translation delay (τ_TxTl_) of HES7 was estimated to be longer in human PSM (30 ± 1 min) as compared with mouse PSM (17 ± 2 min) (Fig. 3f, top). The fitting also allowed us to estimate the degradation rate of HES7 mRNA (δ_m_), showing slower degradation in human PSM (half-life in mouse: 10 ± 0.3 min; in human: 16 ± 0.3 min) (Fig. 3f, bottom). Note that the HES7 gene used in these measurements did not include the introns (see Fig. 3b, d). Since introns affect mRNA splicing and therefore serve as another source of delays^14,30,32^, we measured the delay caused by HES7 introns by creating HES7 promoter-luciferase reporters with (w/) and without (w/o) HES7 introns (Fig. 3g, h) and estimating the phase difference between the oscillations of the two reporters (Fig. 3g; Supplementary Fig. 7). Again, the HES7 intron delay (τ_In_) was longer in human PSM (37 ± 3 min) compared with mouse PSM (13 ± 3 min) (Fig. 3i). Roughly consistent with our measurements, the intron delay or splicing delay in mouse embryos was previously reported to be 12-19 min^14,32^. Finally, to measure the delay for HES7 to start repressing its own promoter, we induced the expression of HES7 and estimated the onset of decline in the HES7 promoter activity (Fig. 3j; Supplementary Fig. 8). Fitting the results to an open loop repression model in which the induced HES7 protein represses the expression of HES7 promoter-luciferase reporter showed that the HES7 repression delay (τ_Rp_) is negligible in both murine and human PSM.

To confirm that the degradation rates and delays measured in both murine and human PSM can indeed explain the interspecies period difference in the segmentation clock, we built a simple mathematical model of HES7 oscillation^20^ based directly on the following parameters: δ_p_, δ_m_, τ_TxTl_, τ_In_, and τ_Rp_ (Fig. 3k; see Methods). Note that our mathematical analyses of the model showed that the oscillation period depends on these measured parameters (i.e., degradation rates and total delays), and that other parameters, such as the transcription and translation rates and the repression threshold, essentially do not affect the period (Supplementary Text 1)^20^. Even though one unmeasured parameter, the repression Hill coefficient, potentially affects the oscillation period (Supplementary Text 1), varying this parameter within a realistic range did not change the period dramatically (Fig. 3l). Remarkably, our simulation of oscillations with the murine parameters showed periods of ~150 min whereas that with human parameters showed ~300 min periods (Fig. 3l), reproducing the 2-3-fold period difference between actual murine and human PSM (see Fig. 1e). These results mean that the slower biochemical reactions of HES7 (i.e., slower degradations and longer delays) in human PSM as compared with those in mouse PSM are sufficient to quantitatively account for the longer oscillation period of the human segmentation clock.

Next, we explored how universal our finding of slower biochemical reactions in human cells is. To test whether it is specific to the HES7 gene or generalizable to other genes, we measured the degradation rates of six other genes, transcription factors expressed at the PSM stage^7^ (Fig. 4a, b; Supplementary Fig. 9). GBX2, MSGN1, and TBX6 proteins showed slower degradation rates in human PSM than in mouse PSM, whereas CDX2, EVX1, and Brachyury T did not (Fig. 4c). We also measured the transcription and intron delays (τ_Tx_, τ_In_) (Fig. 4d, e; Supplementary Fig. 10). TBX6, GBX2, and MSGN1 showed longer delays in human PSM than in mouse PSM whereas EVX1 did not show a significant interspecies difference (Fig. 4f). These results suggest that the slower biochemical reactions in human PSM with respect to mouse PSM can extend to several other genes, but not to all genes.

**Figure 4.**
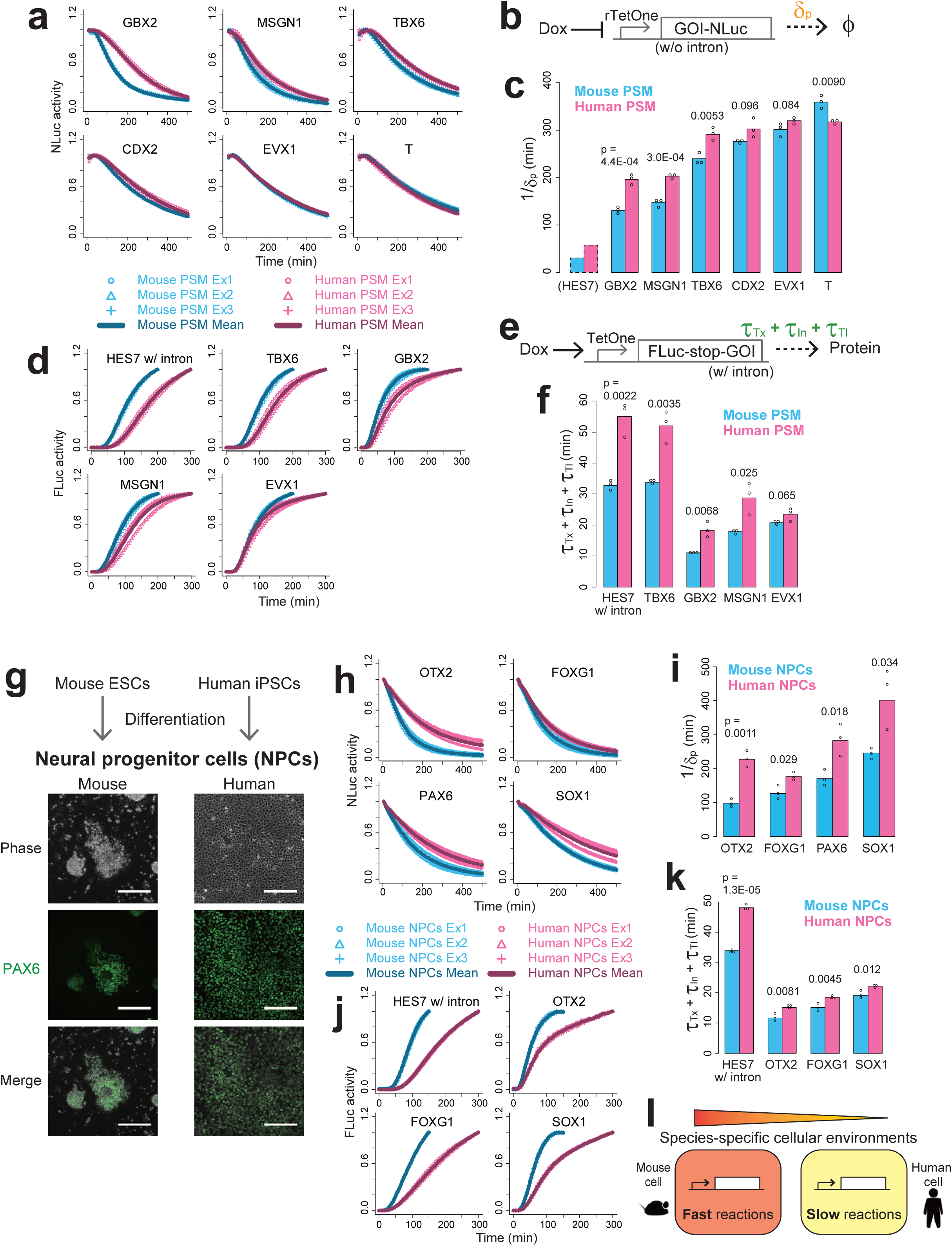
Generality of slower biochemical reactions in human cells. **a, b**, Degradation assay of other genes expressed at the PSM stage. The transcription of a gene of interest (GOI) fused with NLuc was halted upon Dox addition at t = 0 (b), and the decay of NLuc signal was monitored in either murine or human PSM (a). **c**, Inverse protein degradation rates of other PSM marker genes estimated from Supplementary Fig. 9. The HES7 degradation rate shown in Fig. 3c is displayed as a control. **d, e**, Expression delay assay of other genes expressed at the PSM stage. The transcription of FLuc-GOI fusion construct flanked by a stop codon was induced upon Dox addition at t = 0 (e), and the onset of FLuc signal was monitored in either murine or human PSM (d). Note that the delay measured here is the sum of the transcription delay of the fusion construct, intron delay of GOI, and translation delay of FLuc. Brachyury T and CDX2 were not used due to their long introns. **f**, Delays of other PSM genes estimated from Supplementary Fig. 10. **g**, *In vitro* differentiation of NPCs from mouse ESCs and human iPSCs. PAX6 is a neural marker gene. Scale bars: 200 μm. **h**, Degradation assay in NPCs. The transcription of GOI-NLuc was halted upon Dox addition at t = 0, and the decay of NLuc signal was monitored in either murine or human NPCs. **i**, Inverse protein degradation rates in NPCs estimated from Supplementary Fig. 11. **j**, Expression delay assay in NPCs. The transcription of FLuc-GOI fusion construct flanked by a stop codon was induced upon Dox addition at t = 0, and the onset of FLuc signal was monitored in either murine or human NPCs. PAX6 was not used due to its long introns, and HES7 in NPCs was used as a control. **k**, Delays in NPCs estimated from Supplementary Fig. 12. **l**, Proposed scheme. Murine and human cells have different cellular environments that affect the speeds of several biochemical reactions. All p-values are from two-sided student’s t-test.

Finally, to test whether the slower biochemical reactions in human cells are specific to PSM or generalizable to other cell types, we induced neural progenitor cells (NPCs) from mouse ESCs and human iPSCs (Fig. 4g)^33,34^. We measured the degradation rates of neural marker genes^34^ in both murine and human induced NPCs (Fig. 4h; Supplementary Fig. 11). All the four proteins tested, OTX2, FOXG1, PAX6, and SOX1, showed slower degradation rates in human NPCs as compared with mouse NPCs (Fig. 4i). We also measured the transcription and intron delays of OTX2, FOXG1, and SOX1 (Fig. 4j; Supplementary Fig. 12), demonstrating slightly longer delays in human NPCs for all three genes (Fig. 4k). These results suggest that slower biochemical reactions in human cells can be applicable not only to the PSM fate but also to other cell types, even though more systematic measurements will be necessary in the future. We propose that murine and human cells possess species-specific cellular environments that affect speeds of several biochemical reactions including degradations and delays (Fig. 4l), potentially causing other interspecies differences in developmental time. The cellular environments can mean any gene set or any cellular characteristic, such as the metabolic rate and cell size.

In summary, we have shown that the human segmentation clock exhibits 2-3-times slower oscillations in comparison with mouse, because of slower degradation rates of HES7 and longer delays in its expression in a human PSM cell. An obvious next challenge is to reveal the mechanism by which human cells display slower biochemical reactions. Since our results have revealed the existence of several other genes that show different reaction speeds between murine and human cells, it may be interesting to classify the genes that show such an interspecies difference and to find commonalities among them. Another future challenge is to investigate developmental time of other species than mouse and human. Interestingly, delays due to splicing and export of mRNAs of HES-related genes have previously been reported to be different among mouse, chick, and zebrafish^32^. Because ESCs and iPSCs of diverse mammals are now available^35,36^, their *in vitro* differentiation will enable to compare the same cell types among different mammalian species and to study their different tempos of development.

## Methods

### Pluripotent stem cell cultures

Mouse ESCs (EB5, a gift from H. Niwa) were maintained on gelatin coated dish with GMEM containing 10% KSR, 1% FBS, nonessential amino acids (1 mM), β-mercaptoethanol (0.1 mM), sodium pyruvate (1 mM), LIF (2000 U/ml), CHIR99021 (3 µM), and PD0325901 (1 µM). Human iPSCs (201B7, feederless) were maintained on iMatrix-511 silk (Nippi) coated dishes or plates with StemFit AK02N medium (Ajinomoto).

### DNA constructs

The genetic constructs are listed in Supplementary Table 2. The promoters or genes were subcloned into pDONR vector to create entry clones. These entry clones were recombined with *piggyBac* vector (a gift from K. Woltjen)^37^ by using the Multisite Gateway technology (Invitrogen). The DNA constructs were introduced into the cells with Amaxa Nucleofector (Lonza).

### Induction of murine and human PSM

Mouse ESCs were first cultured in N2B27 medium containing 1% KSR, ACTIVIN A (20 ng/ml), and bFGF (10 ng/ml) for 4 days and converted to mouse EpiLCs^38,39^. The induced mouse EpiLCs were further cultured in CDMi^40^ containing SB431542 (10 μM), DMH1 (2 μM), CHIR99021 (10 μM), and bFGF (20 ng/ml) for 2 days to induce mouse PSM cells. To induce human PSM cells, our 1 step induction protocol^9^ was mainly used. Human iPSCs were seeded on a 35 mm dishes coated with iMatrix-511 silk or matrigel and cultured for 4 days. Then the cells were cultured in CDMi containing SB431542 (10 μM), DMH1 (2 μM), CHIR99021 (10 μM), and bFGF (20 ng/ml) for another 3.5 days to induce human PSM cells. For the degradation assay, our 2 step induction protocol^12^ was used. Human iPSCs were seeded on a 35 mm dish coated with iMatrix-511 silk and cultured for 5 days. Then the medium was changed into CDMi containing bFGF (20 ng/ml), CHIR99021 (10 μM), and ACTIVIN A (50 ng/ml) for 24 hours to induce primitive streak (PS) cells. The induced PS cells were further cultured in CDMi containing SB431542 (10 μM), CHIR99021 (3 μM), LDN-193189 (250 nM), and bFGF (20 ng/ml) for 24 hours to induce human PSM cells.

### Induction of murine and human NPCs

To induce mouse NPCs, mouse ESCs were seeded on a gelatin coated dish and cultured in the NDiff 227 medium (TAKARA) for 5 days^33^. To induce human NPCs, human iPSCs were seeded on a matrigel coated dish and cultured in the STEMdiff SMADi Neural Induction medium (STEMCELL Technologies) for 7 days^34^. NPC differentiation was checked by immunostaining with an anti-PAX6 antibody (BioLegend).

### Oscillation analyses

After the induction of murine or human PSM cells, the medium was changed into CDMi containing SB431542 (10 μM), DMH1 (2 μM), CHIR99021 (1 μM), bFGF (20 ng/ml), and D-luciferin (200 μM or 1mM) to monitor oscillations of the HES7 reporter signal. For the single cell imaging, the induced PSM cells were re-seeded on iMatrix-511 silk coated dish. After 6 hours, the medium was changed into CDMi containing SB431542 (10 μM), DMH1 (2 μM), CHIR99021 (1 μM), bFGF (20 ng/ml), Latrunculin A (0.5 μM)^41^, and D-luciferin (1 mM). Bioluminescence was measured with Kronos Dio Luminometer (Atto) or LCV110 microscope (Olympus). The obtained signal was detrended by using a 60 min (mouse), 90 min (human), or 100 min (human single cell) moving average-subtraction method. The data displayed were normalized to the 1st peak of oscillations. The oscillation period was defined as the time interval between the 1st and 4th peaks divided by 3 cycles. For *ex vivo* measurements, the period was defined with the 1st and 3rd peaks.

### HES7 loci swapping

The CRISPR guide sequences for HES7 swapping were cloned into pSpCas9(BB)-2A-GFP vector (addgene #48138)^42^ (see Supplementary Table 2). As the template for homologous recombination, a bacterial artificial chromosome (BAC) containing human HES7 locus (RP11-769H22)^43^ was obtained from BACPAC resources center (Children’s Hospital & Research Center at Oakland). After homology arms were inserted (see Supplementary Table 2), the purified BAC was introduced into mouse ESCs with CIRSPR/Cas9 guides to swap the HES7 loci between mouse and human.

### Southern blotting

Southern blotting was performed according to the DIG Application Manual for Filter Hybridization (Roche) using the PCR DIG Probe Synthesis kit (Roche). The probe sequences are available in Supplementary Table 2.

### HES7 knockout

The CRISPR guide sequences for HES7 knockout were cloned into pSpCas9(BB)-2A-GFP (see Supplementary Table 2). HES7 knockout was performed using transient transfection of multiple CRISPR guide constructs. The deletion of HES7 was confirmed by PCR.

### Transgenic mice and *ex vivo* tissue culture

HES7 swapping was performed in mouse ESCs (TT2)^44^. Two hetero swap ESC clones were isolated, and chimeric mice were generated according to standard procedures. The body segments of transgenic mice were imaged at 4 weeks with *in vivo* micro X-ray CT System R_mCT (RIGAKU). Mice carrying the HES7 reporter pH7-UbLuc-In (−) were previously described^14^. Time-lapse imaging of *ex vivo* culture was performed as described previously^13^ with several modifications. Mouse embryos were collected at 10.5 dpc and dissected in PBS containing 0.2% BSA. Tail portions from Wt and Homo swap embryos were embedded in 0.2% low-melting point agarose in a silicon ring set onto a 35 mm glass-bottom dish, and then cultured in DMEM/F12 containing 1% BSA, L-Glutamine (2 mM), D-Glucose (1 g/L), HEPES (15 mM), and D-luciferin (1 mM) with 5% CO2 and 80% O2. Imaging was performed on IX81 microscope (Olympus) equipped with VersArray CCD camera (Princeton Instruments). All animal experiments were approved by the Institutional Animal Care and Use Committee of RIKEN Kobe Branch or Kyoto University, and performed according to animal experimentation guidelines of RIKEN and Kyoto University.

### Models and parameter measurements

#### 1. HES7 oscillation model

To simulate the oscillation of HES7, previously proposed delay differential equations of HES feedback loop were used^20^.

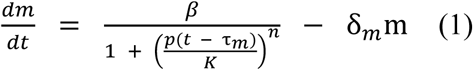

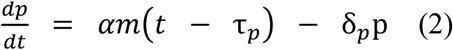

where m and p are the concentrations of mRNA and protein, respectively. δ_m_ and δ_p_ are the degradation rates of mRNA and protein, α and β are the translation and transcription rates, K is the repression threshold, and n is the repression Hill coefficient. τ_m_ and τ_p_ are the mRNA and protein delays, and they have the following relationships with the experimentally measured delays:

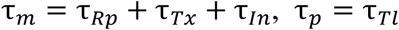

where τ_Rp_, τ_Tx_, τ_In_, and τ_Tl_ are the repression, transcription, intron, and translation delays, respectively. The parameter values are summarized in Supplementary Table 1. The numerical calculation was performed with dde23 of MATLAB (Mathworks), and the oscillation periods were estimated by computing the power spectra of the time series.

#### 2. Degradation assay of HES7

The overexpression of a fusion construct of HES7 and NLuc was regulated by the rTetOne system (reverse TetOne system; see Supplementary Table 2). The construct was introduced into mouse ESCs or human iPSCs where the endogenous HES7 was knocked out. After PSM cells were induced in the presence of Doxycycline (Dox; 100 ng/m), the expression of the fusion protein was initiated by washing out Dox and changing the medium into CDMi containing protected furimazine (Promega; 1 μM). After the NLuc signal was confirmed 5-8 hours later, the expression of the fusion protein was halted by Dox (300 ng/ml) addition, and the decay of NLuc signal was monitored with Kronos Dio luminometer. To exclude the influence of residual mRNAs, only the later time points where the NLuc signal displayed a single exponential decay curve were used. To estimate the protein degradation rate (δ_p_) of HES7, the slope of log-transformed data was calculated with the least square method of R.

#### 3. Expression delay assay of HES7

The overexpression of a fusion construct of HES7 (w/o intron) and NLuc was regulated by the TetOne system. The construct was introduced into mouse ESCs or human iPSCs where the endogenous HES7 was knocked out. After PSM cells were induced in the absence of Dox, the medium was changed into CDMi containing protected furimazine (1μM). Six hours after the medium change, the expression of the fusion protein was initiated by Dox (300 ng/ml), and the onset of NLuc signal was monitored with Kronos Dio luminometer. To estimate the sum of the transcription delay and translation delay (τ_TxTl_) of HES7 as well as the mRNA degradation rate (δ_m_) of HES7, models for the expression delay assay and degradation assay were constructed.

Expression delay model:

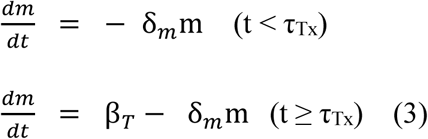

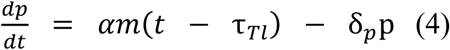

where β_T_ is the transcription rate of the TetOne promoter.

The solution of this is

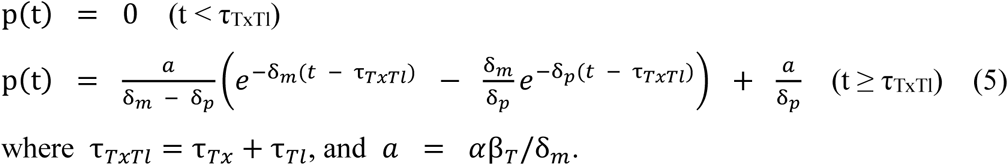

where τ_*TxTl*_ = τ_*Tx*_ + τ_*Tl*_, and α = αβ_*T*_/δ_*m*_.

Degradation model:

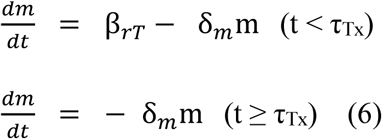

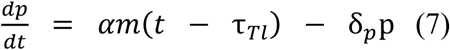

where β_rT_ is the transcription rate of the rTetOne promoter.

The solution of this is

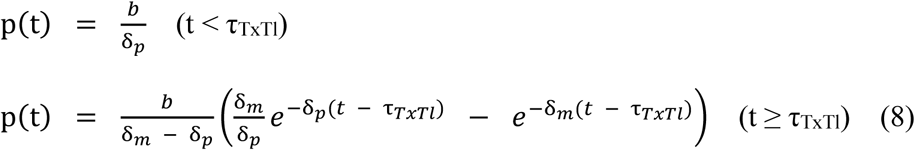

where *b* = αβ_*rT*_/δ_*m*_.

As for δ_p_, the value estimated in the degradation assay was used. τ_TxTl_ and δ_m_, together with a and b, were estimated in Python by simultaneously fitting the data of expression delay assay and degradation assay to the equations (5) and (8), respectively, with SciPy’s basin-hopping algorithm.

#### 4. Degradation and expression delay assays of other genes

For the degradation assay, the overexpression of a fusion construct of a target gene and NLuc was regulated by the rTetOne system. The construct was introduced into mouse ESCs or human iPSCs. After PSM cells or NPCs were induced in the presence of Dox (100 ng/m), the expression of the fusion protein was initiated by washing out Dox and changing the medium into CDMi containing protected furimazine (5 μM in human NPCs, 1 μM in the other cell types). After the NLuc signal was confirmed 5-8 hours later, the expression of the fusion protein was halted by Dox (300 ng/ml), and the decay of NLuc signal was monitored with Kronos Dio luminometer. To estimate the degradation rate (δ_p_) of the fusion protein, the slope of log-transformed data was calculated with the least square method of R.

For the expression delay assay, the overexpression of a fusion construct of FLuc (w/ stop codon) and a target gene (w/ intron) was regulated by the TetOne system. The construct was introduced into mouse ESCs or human iPSCs. After PSM cells or NPCs were induced in the absence of Dox, the medium was changed into CDMi containing D-luciferin (200 μM). Six hours after the medium change, the expression of the fusion construct was initiated by Dox (300 ng/ml), and the onset of FLuc signal was monitored with Kronos Dio luminometer. To estimate the sum of the transcription delay, intron delay, and translation delay (τ_TxInTl_) of the fusion construct, a model for expression delay assay was constructed.

Expression delay model:

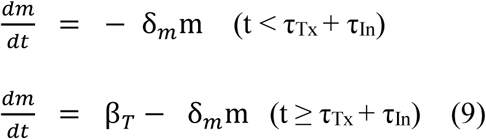

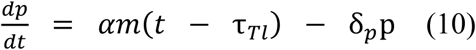

The solution of this is

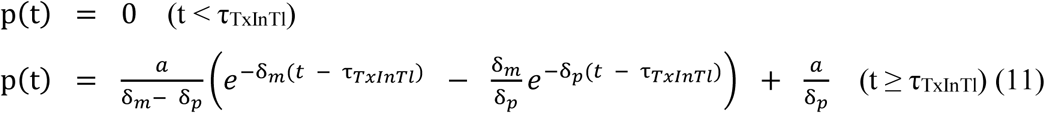

where τ_*TxInTl*_ = τ_*Tx*_ + τ_*In*_ + τ_*Tl*_.

δ_TxInTl_ for each target gene, together with δ_m_, δ_p_, and a, were estimated in Python by fitting the data of expression delay assay to the equation (11) with SciPy’s basin-hopping algorithm.

#### 5. Intron delay assay

The HES7 promoter-NLuc-stop-HES7 (w/o intron) and HES7 promoter-FLuc-stop-HES7 (w/ intron) reporter constructs were introduced into mouse ESCs or human iPSCs. After PSM cells were induced, the medium was changed into CDMi containing protected furimazine (1 μM) and D-luciferin (1 mM), and the oscillations of the NLuc and FLuc signals were simultaneously monitored with Kronos Dio luminometer. To estimate the intron delay (τ_In_) of HES7, the oscillation phase difference between the ‘w/o intron’ and ‘w/ intron’ reporters was estimated by calculating their cross correlation with R. To normalize the difference in the maturation/degradation time between NLuc and FLuc, cells containing the HES7 promoter-NLuc-stop-HES7 (w/o intron) and HES7 promoter-FLuc-stop-HES7 (w/o intron) constructs were also created, and the phase difference between the NLuc and FLuc reporters was subtracted from that between the w/o intron and w/ intron reporters.

#### 6. Repression delay assay

First, the expression delay assay of FLuc reporter was performed to estimate the degradation rates of the mRNA (δ_f_) and protein (δ_F_) of FLuc as well as the transcription/translation delay (τ_TxTlF_) of FLuc as described in the section of Expression delay assay of other genes. Next, the overexpression of a fusion construct of HES7 (w/o intron) and NLuc was regulated by the TetOne system. The expression of HES7 promoter-FLuc reporter was repressed by the HES7-NLuc protein. The constructs were introduced into mouse ESCs or human iPSCs where the endogenous HES7 was knocked out. After PSM cells were induced in the absence of Dox, the medium was changed into CDMi containing D-luciferin (200 μM). Six hours after the medium change, the expression of HES7-NLuc protein was initiated by Dox (300 ng/ml), and the onset of decline in the FLuc reporter signal was monitored with Kronos Dio luminometer. To estimate the repression delay (τ_Rp_) of HES7, a model for the repression delay assay was constructed.

Repression delay model:

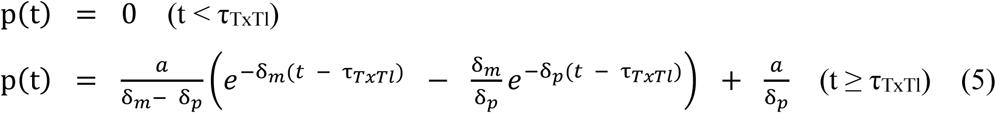

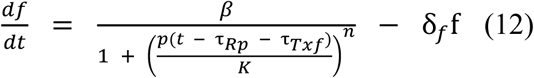

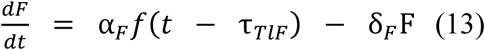

where f and F are the mRNA and protein concentrations of FLuc, respectively. τ_Txf_ and τ_TlF_ are the transcription and translation delays of FLuc (τ_TxTlF_ = τ_Txf_ + τ_TlF_), and α_F_ is the translation rate of FLuc. The numerical calculation was performed with Python, and the resulting F(t) was multiplied by C(t) to incorporate the effect of cell population growth.

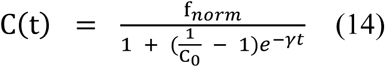

where C_0_ is the initial cell density, γ is the growth rate, and f_norm_ is the scaling factor for luminescence. As for δ_p_, δ_m_, δ_F_, δ_f_, τ_TxTl_, and τ_TxTlF_, measured values were used. The data of repression delay assay were fitted to F(t)×C(t) manually. The fitting was good when τ_Rp_ = 0 with both murine and human parameters.

## Acknowledgments

We thank to J. Sharpe, V. Trivedi, X. Diego, and C. Villava for their comments. We also thank to M. Ogawa and T. Tsuji for helping μCT scan of knock-in mice, to the Laboratory for Animal Resources and Genetic Engineering (LARGE), RIKEN BDR, for generating transgenic mice and animal housing, and to M. Matsumiya for helping oscillation analyses. This work was supported by internal grants from RIKEN and EMBL, Takeda Science Foundation (to M.E.), Grant-in-Aid for Scientific research (KAKENHI) programs from Ministry of Education Culture, Sports, Science, and Technology (MEXT) (16KT0178, 17H05777 to M.M.), and RIKEN Special Postdoctoral Researchers (SPDR) fellowship (to M.M.).

## Author contributions

M.M., M.E., and C.A. conceived the usage of murine and human PSM. M.M. and M.E. conceived the swapping of HES7 loci as well as comparison of biochemical parameters between mouse and human. M.M. and M.E. designed the work and wrote the manuscript. M.M. performed most of the experiments and analyzed the data. H.H. and C.A. developed the induction protocol of mouse PSM. Y.Y., M.I., J.T., and C.A. developed the induction protocol of human PSM. J.G.-O. constructed mathematical models and fitted them to the experimental data. K.Y. and R.K. measured the oscillation period in mouse embryos.

---

**Supplementary Figure 1.**
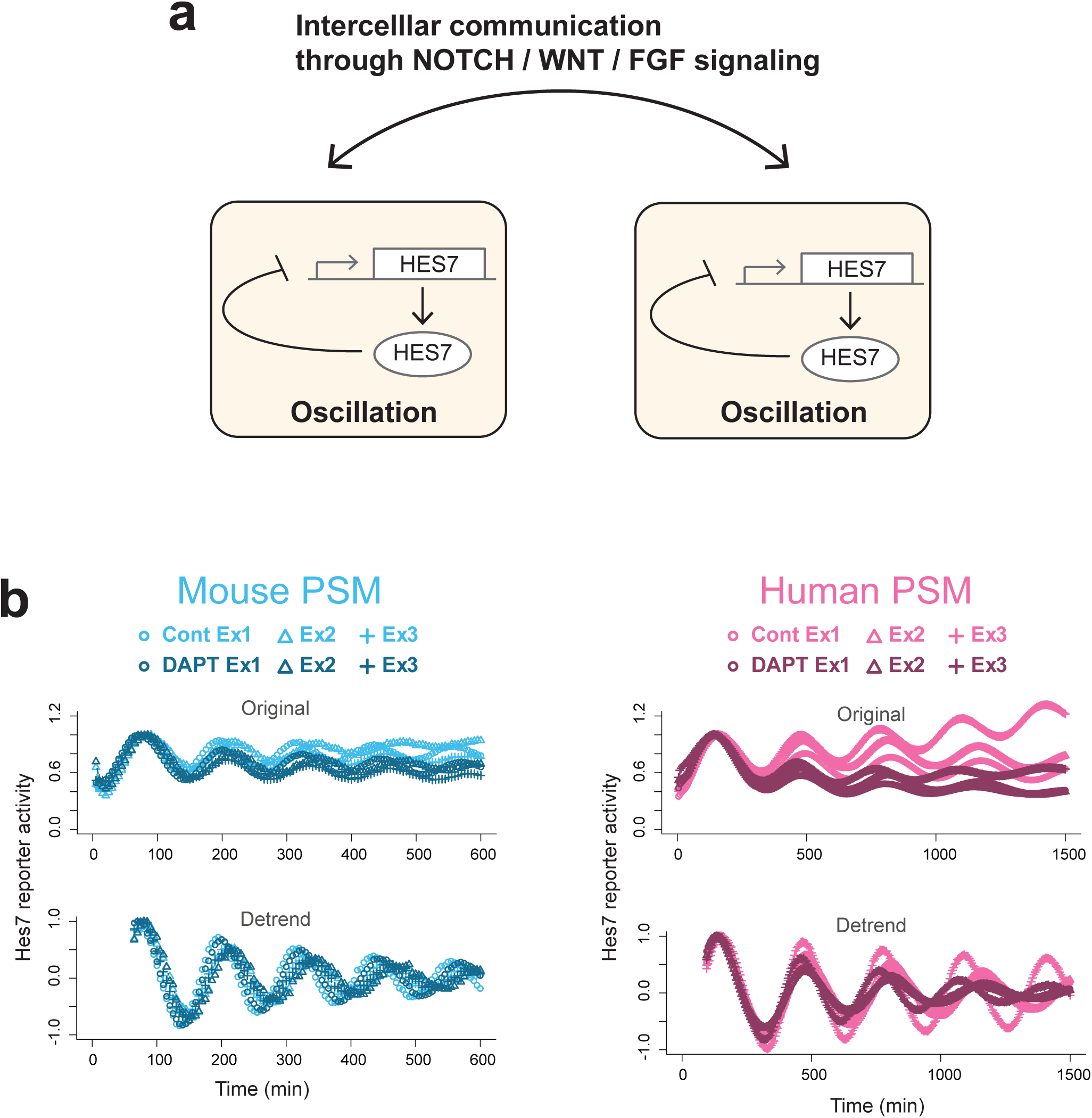
Oscillations and synchronization of the segmentation clock. **a**, Scheme of the segmentation clock. HES7 is a transcription repressor that inhibits its own promoter, giving rise to an oscillatory expression. Oscillations in individual cells are synchronized through intercellular communications driven by NOTCH signaling. WNT and FGF signaling pathways also modulate the segmentation clock. **b**, Effects of inhibiting NOTCH signaling on the HES7 oscillation. The data of Ex2 is also shown in Fig. 1f.

**Supplementary Figure 2.**
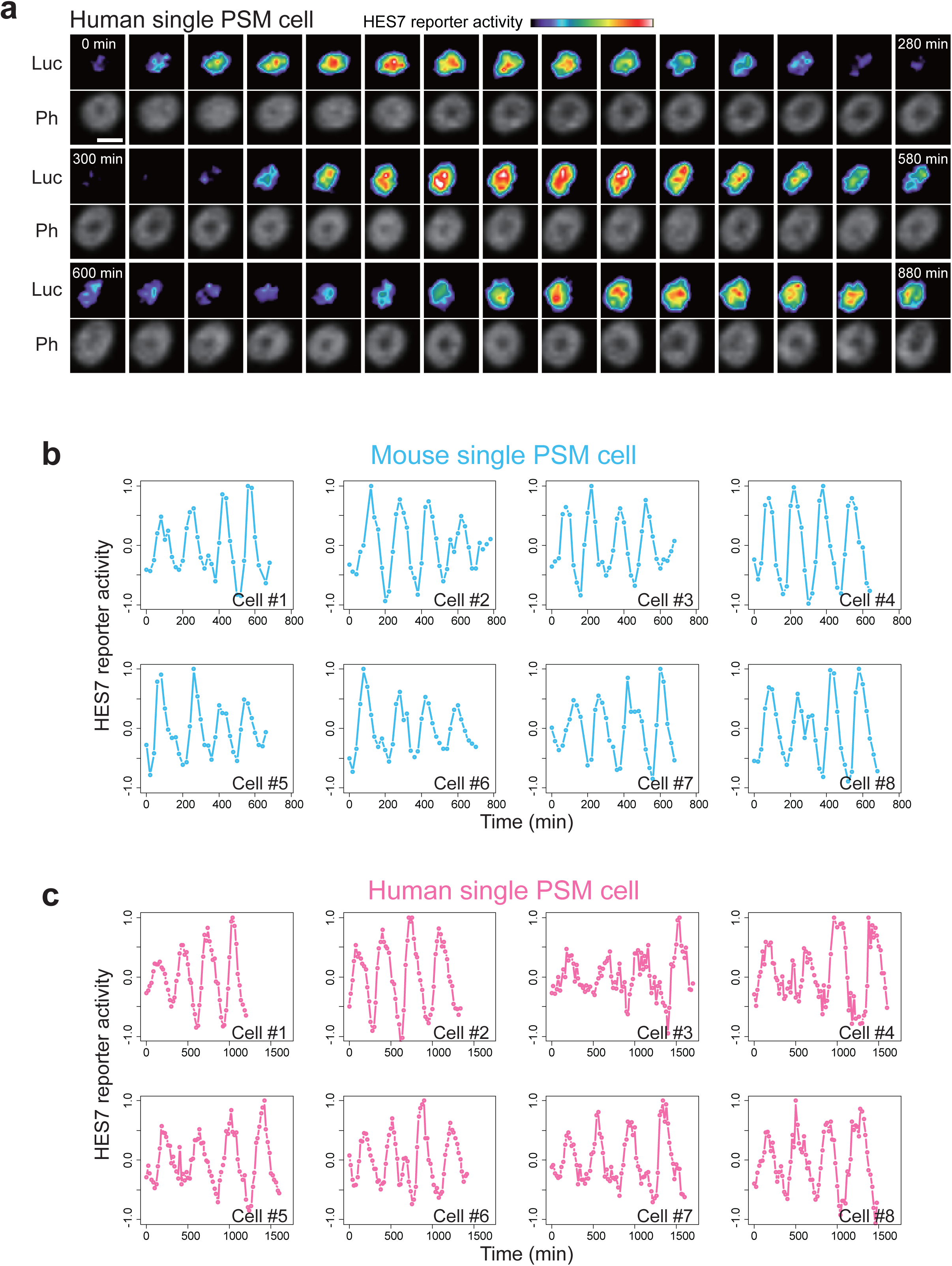
Oscillation in a single cell. **a**, Time-lapse imaging of single cells. Human PSM cells were sparsely split, and the oscillatory HES7 reporter activity in a single cell was monitored. See also Supplementary Video 2. Scale bar: 20 μm. **b**, Oscillations in eight single mouse cells. **c**, Oscillations in eight single human cells.

**Supplementary Figure 3.**
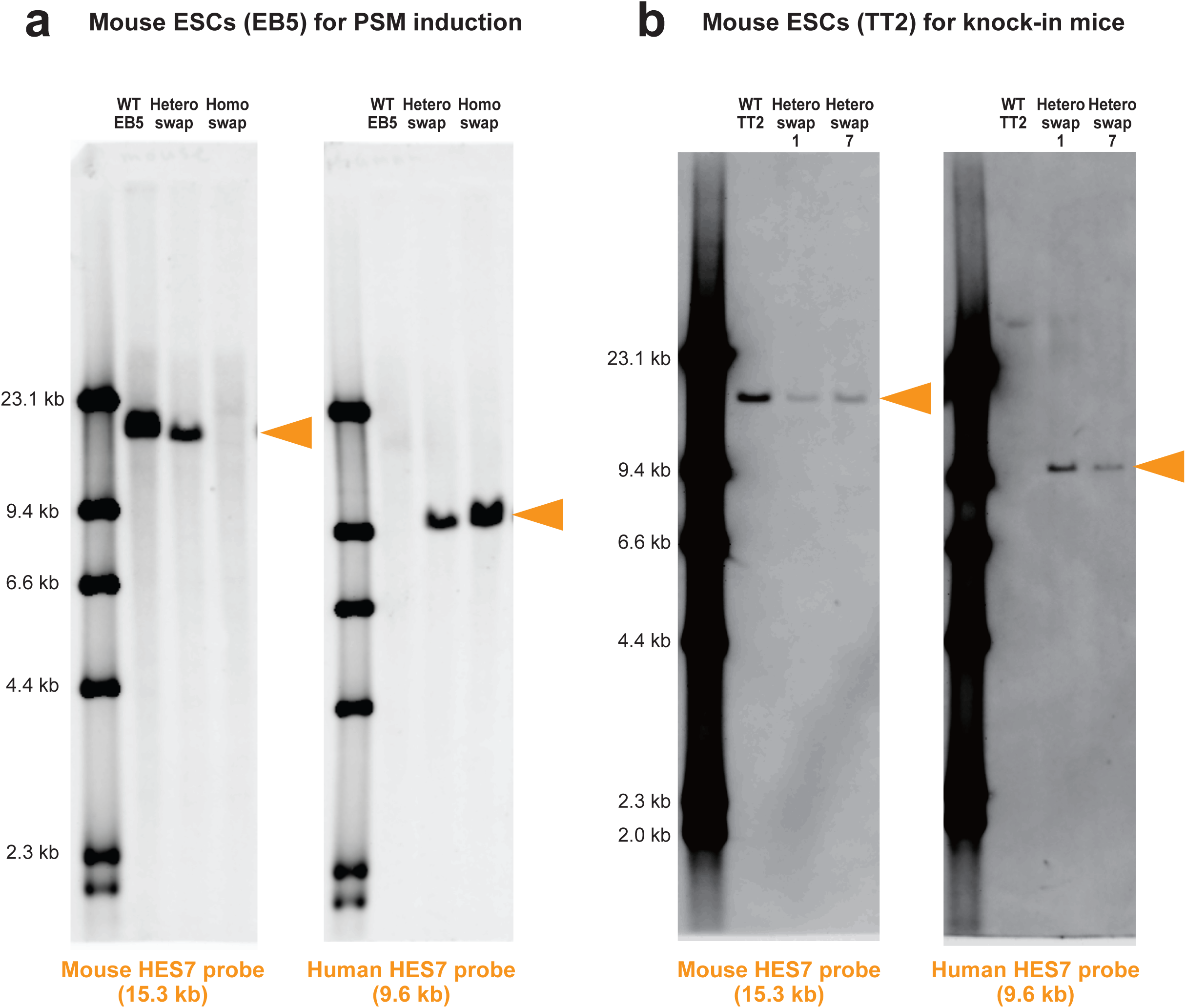
Southern blotting verifying the interspecies genome swapping of the HES7 loci. **a**, Mouse ESCs (EB5) containing the human HES7 locus. The cropped version is shown in Fig. 2b. **b**, Mouse ESCs (TT2) containing the human HES7 locus. Two different lines of hetero swap (clones 1 and 7) were used for knock-in mouse generation.

**Supplementary Figure 4.**
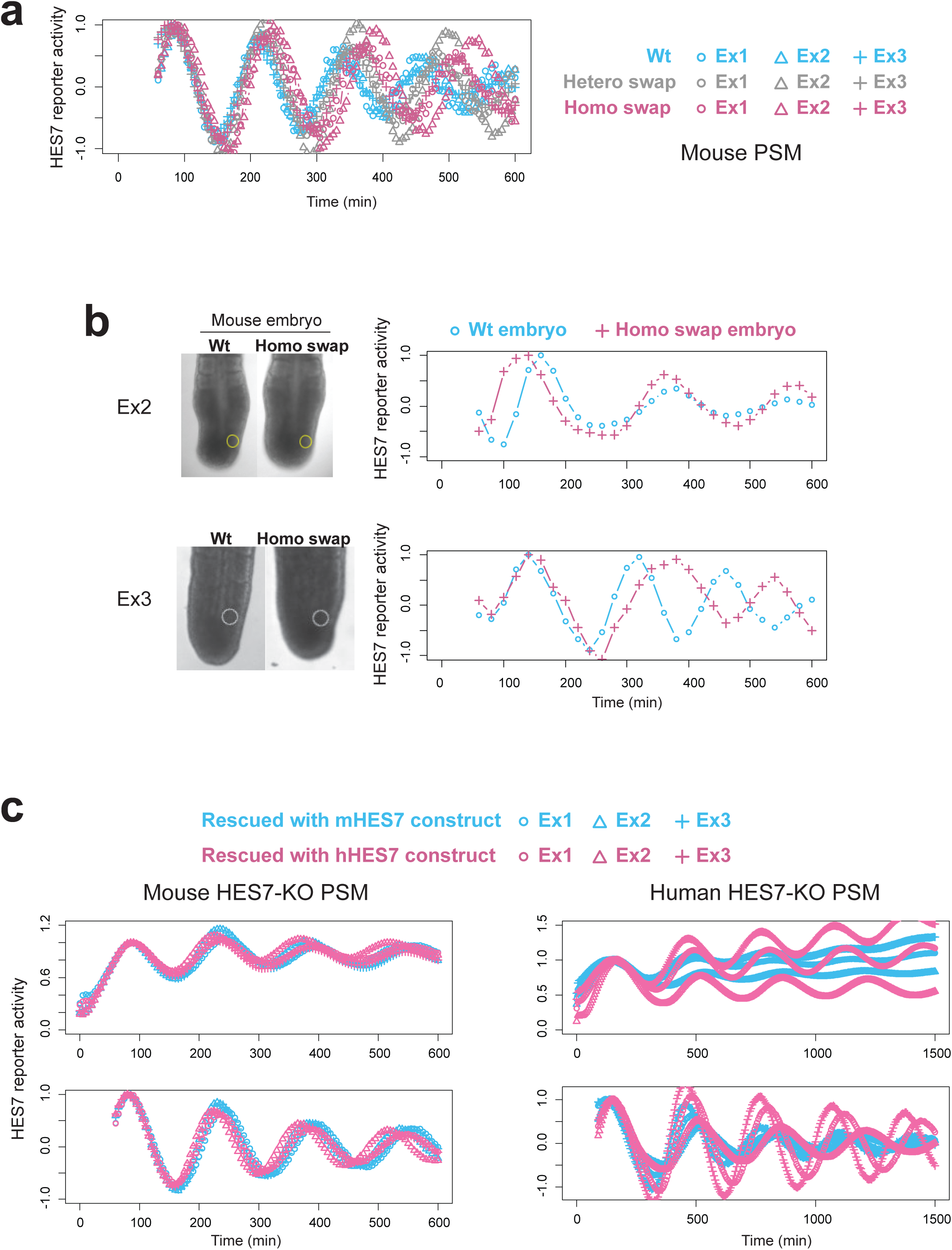
HES7 swapping and KO-and-rescue assay. **a**, Oscillatory HES7 reporter activity in mouse PSM containing the human HES7 locus. Means are shown in Fig. 2c. **b**, *Ex vivo* tail bud cultures of the mouse embryos containing the human HES7 locus. Repeat experiments of Fig. 2f, g. **c**, Rescue of the oscillation by either mouse HES7 or human HES7 construct in HES7-knock-out cells. Means are shown in Fig. 2k, n.

**Supplementary Figure 5.**
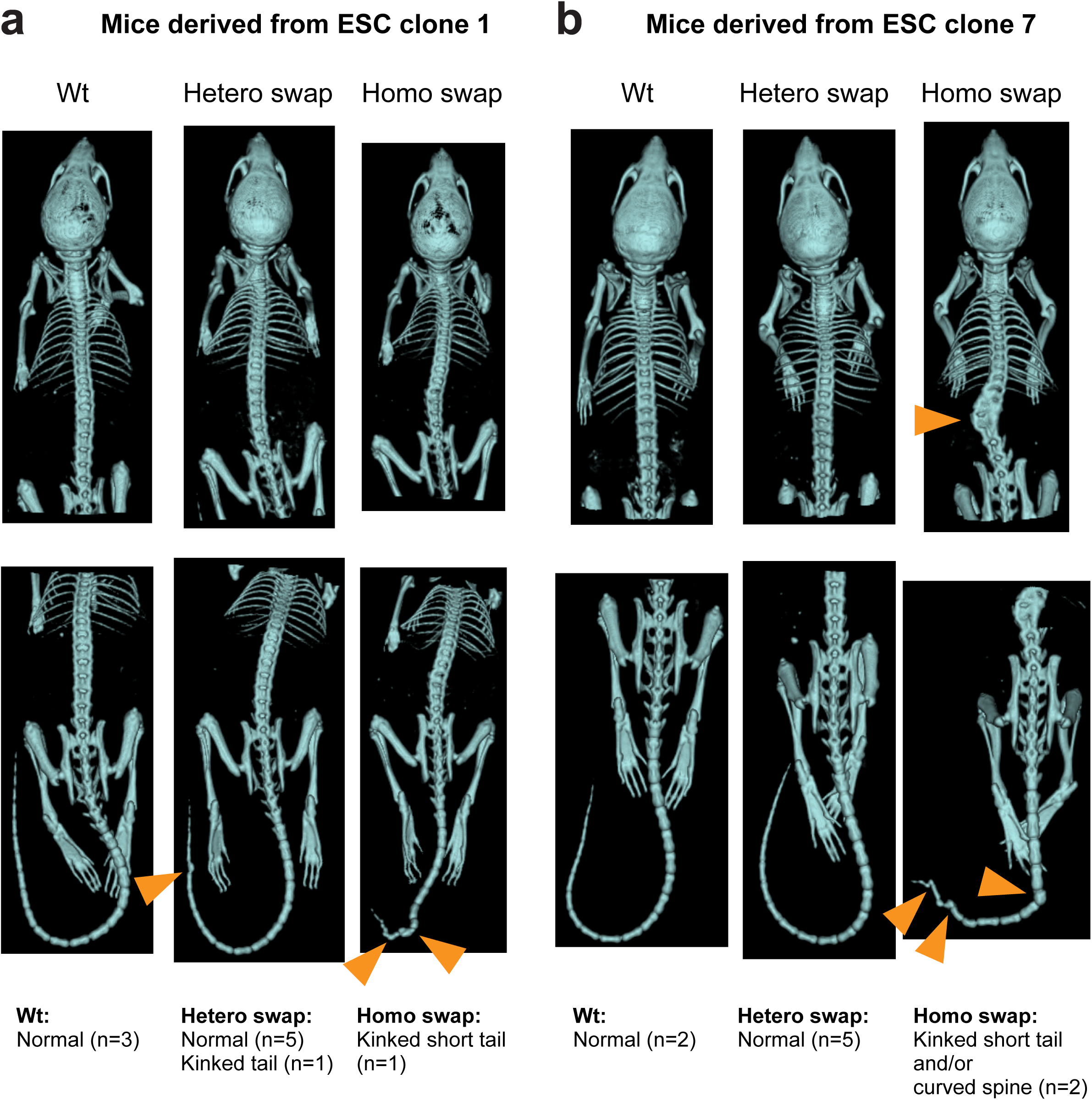
Knock-in mice containing the human HES7 locus. **a**, Phenotypes of swap mice derived from ESC clone 1 shown in Supplementary Fig. 3b. **b**, Phenotypes of swap mice derived from ESC clone 7 shown in Supplementary Fig. 3b. The pictures of clone 7 Wt and Homo swap are also shown in Fig. 2e.

**Supplementary Figure 6.**
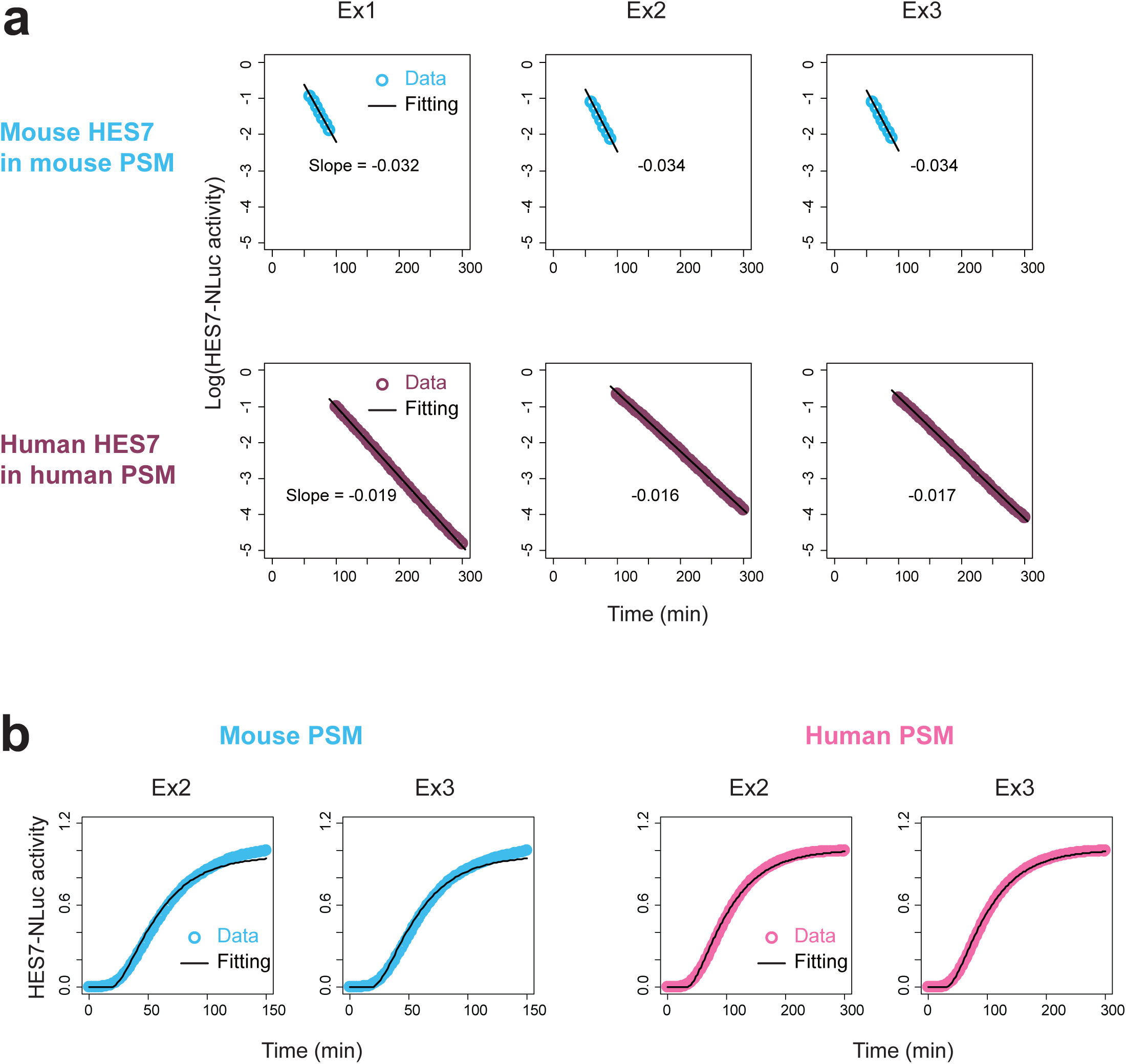
Fitting of the degradation rates and delays of HES7. **a**, Fitting of the degradation rate of HES7 protein. Mouse data at 60-90 min and human data at 100-300 min were used for the fitting. **b**, Fitting of the transcription/translation delay of HES7 shown in Fig. 3d (Ex2, Ex3). Fitting of Ex1 is shown in Fig. 3e. The same data of degradation assay as Fig. 3e was used for fitting.

**Supplementary Figure 7.**
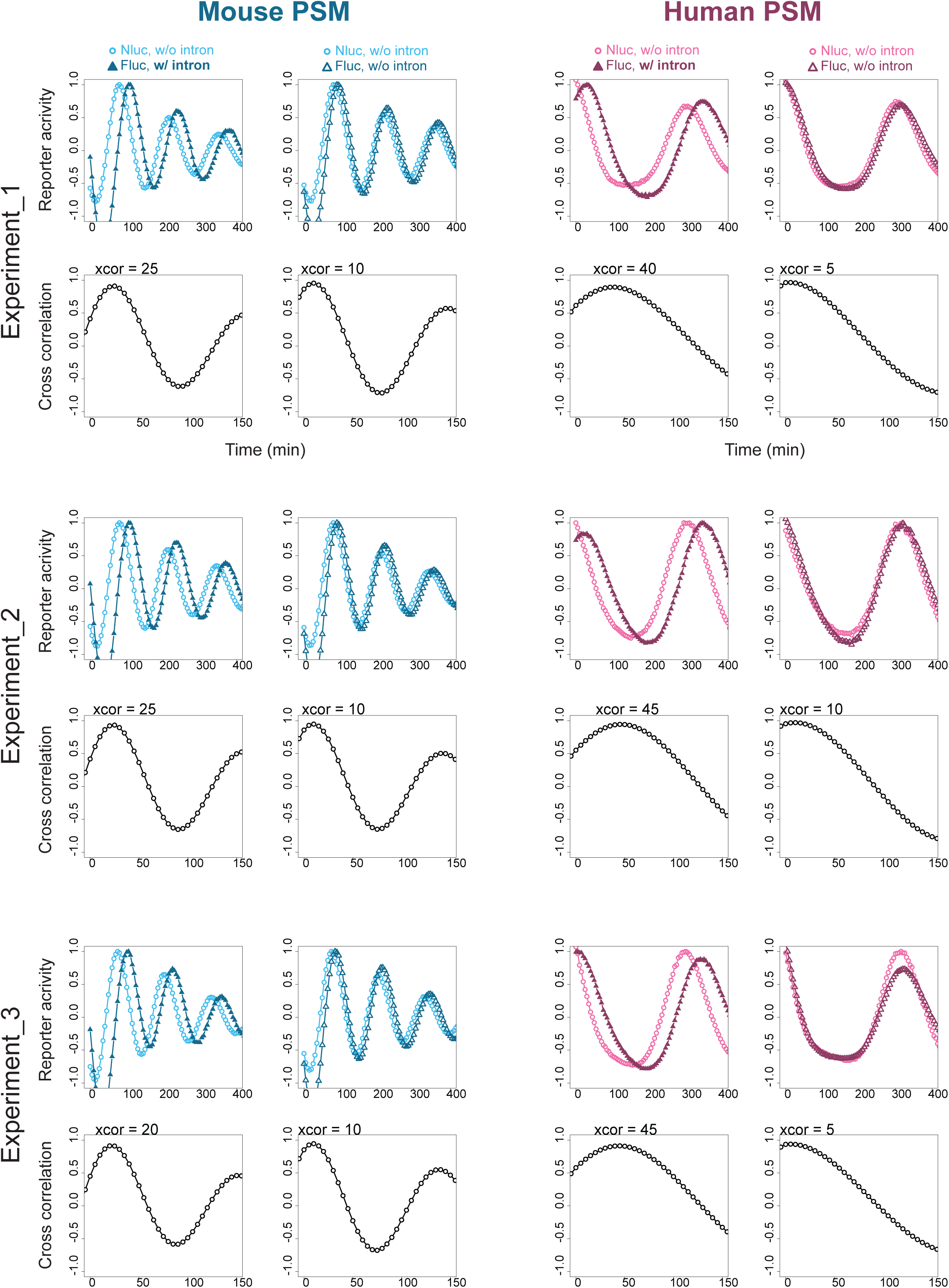
Measurements of the intron delays of HES7. Dual reporter assays and cross correlation functions of NLuc and FLuc signals. Means are shown in Fig. 3g.

**Supplementary Figure 8.**
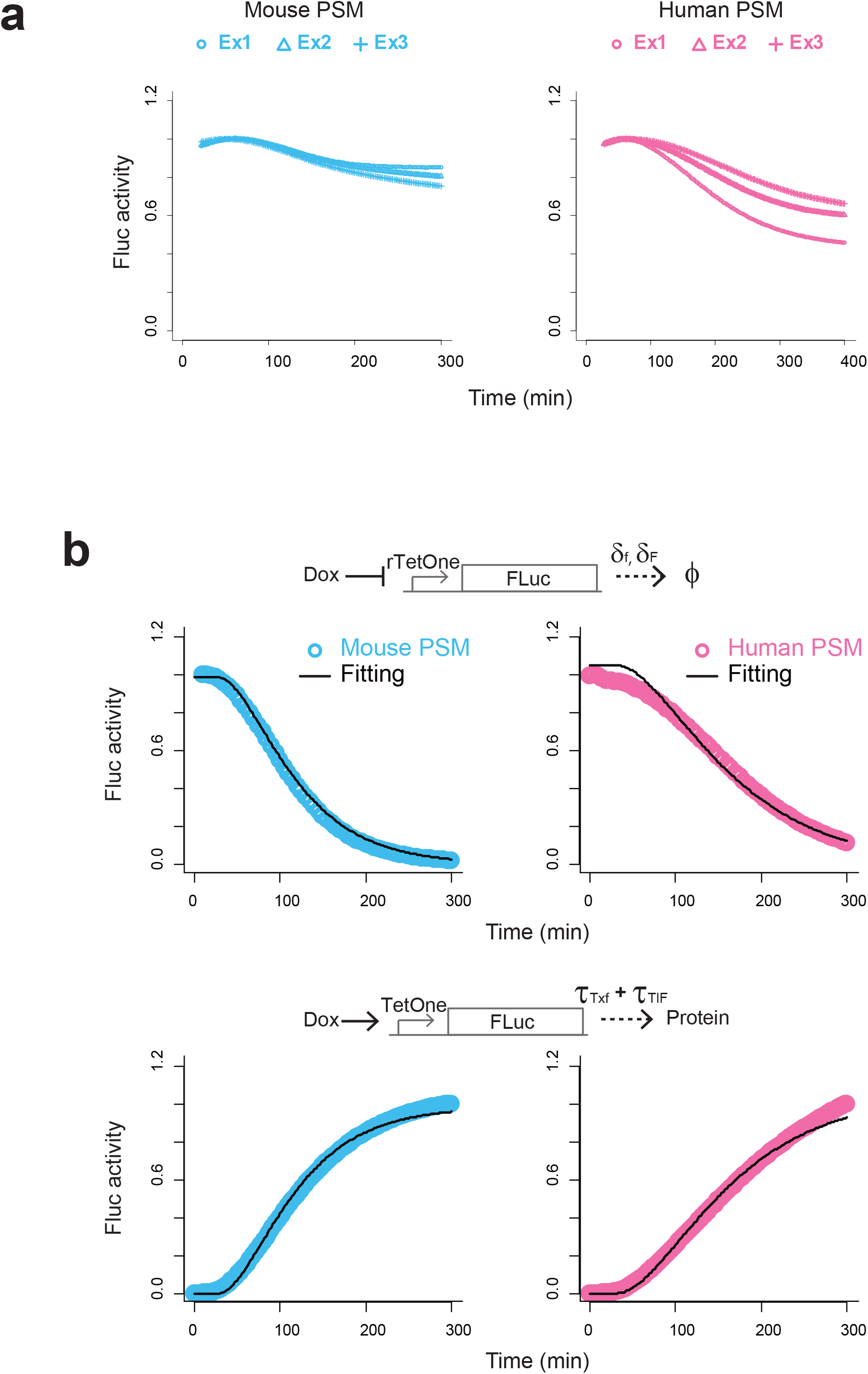
Measurements of the repression delays of HES7. **a**, Measurements of the repression delays of HES7. Means are shown in Fig. 3j. **b**, Measurements and fitting of the degradation rates of mRNA and protein of FLuc (δ_f_, δ_F_; top) and the sum of transcription (τ_Txf_) and translation (τ_TlF_) delays of FLuc (bottom). Mean of three independent experiments. Estimated mouse parameters: δ_f_ = 0.021, δ_F_ = 0.021, τ_TxTlF_ (i.e., τ_Txf_ + τ_TlF_) = 29.3; Estimated human parameters: δ_f_ = 0.014, δ_F_ = 0.014, τ_TxTlF_ = 32.1.

**Supplementary Figure 9.**
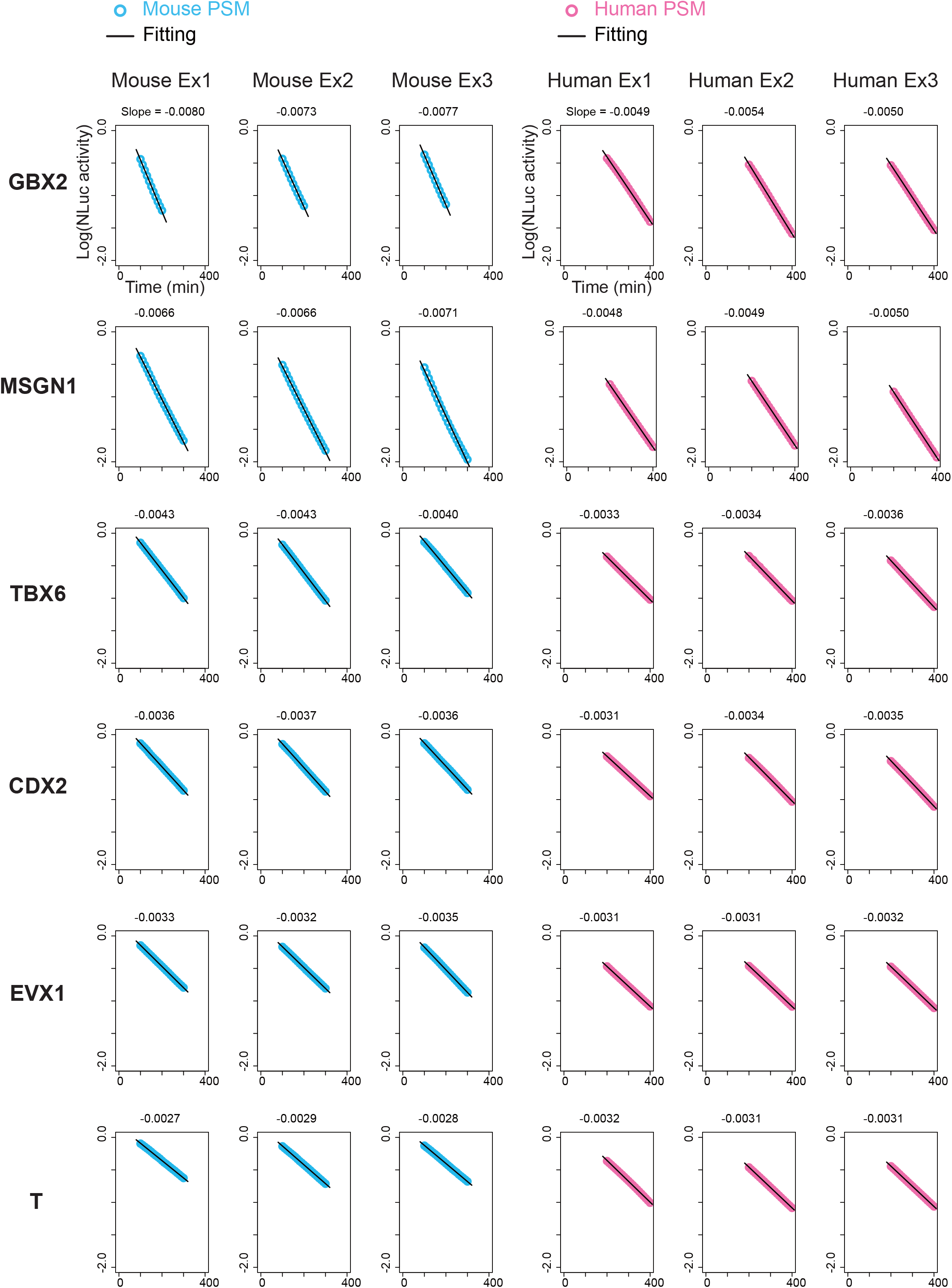
Measurements of the protein degradation rates of other PSM marker genes. Fitting of the protein degradation rates of genes expressed at the PSM stage. Mouse data at 100-300 min (100-200 min for GBX2) and human data at 200-400 min shown in Fig. 4a were used for the fitting.

**Supplementary Figure 10.**
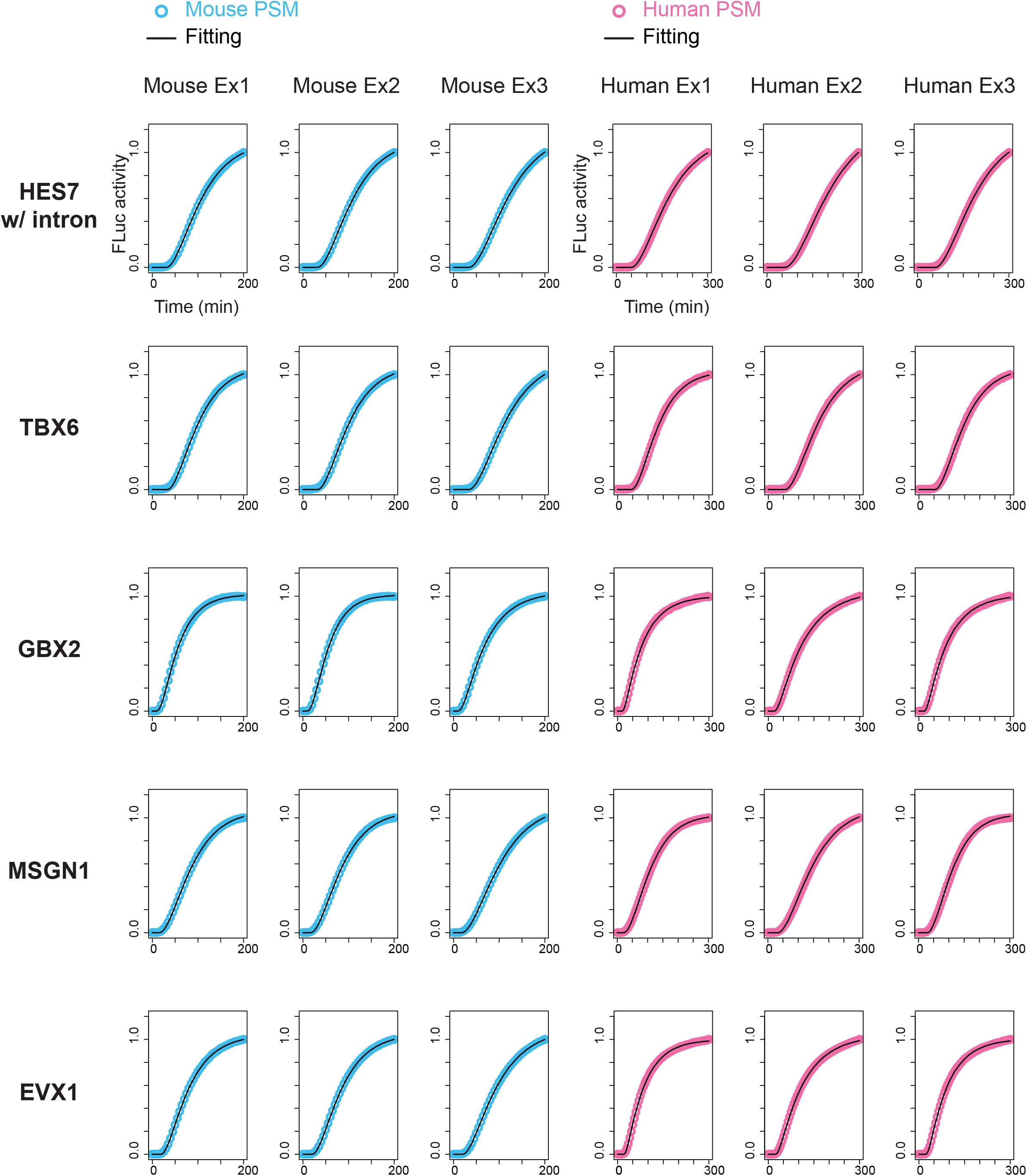
Measurements of the delays of other PSM marker genes. Fitting of the delays of gene expressed at the PSM stage. The original data are shown in Fig. 4d.

**Supplementary Figure 11.**
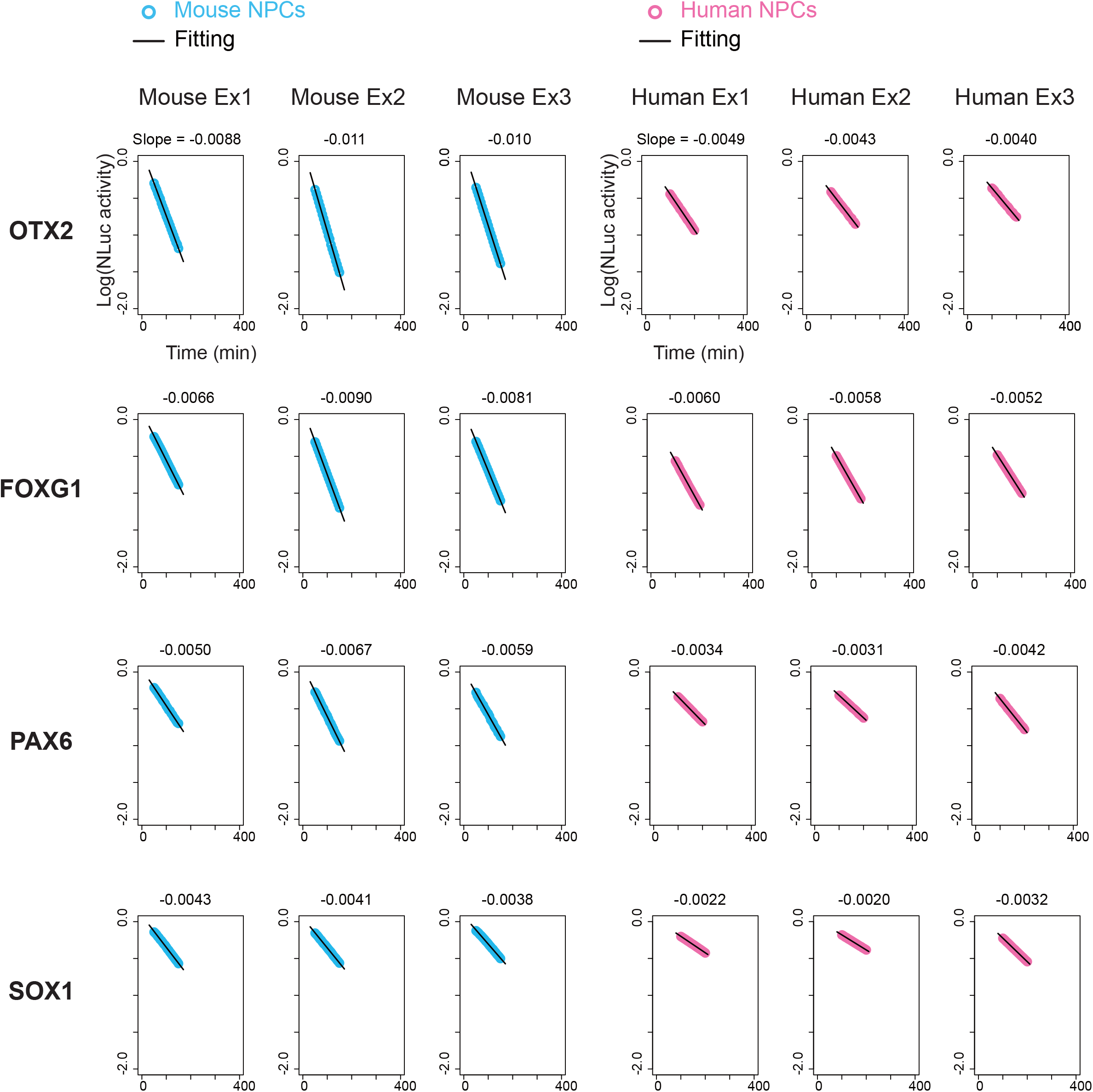
Measurements of the protein degradation rates in NPCs. Fitting of the protein degradation rates in NPCs. Mouse data at 50-150 min and human data at 100-200 min shown in Fig. 4 h were used for the fitting.

**Supplementary Figure 12.**
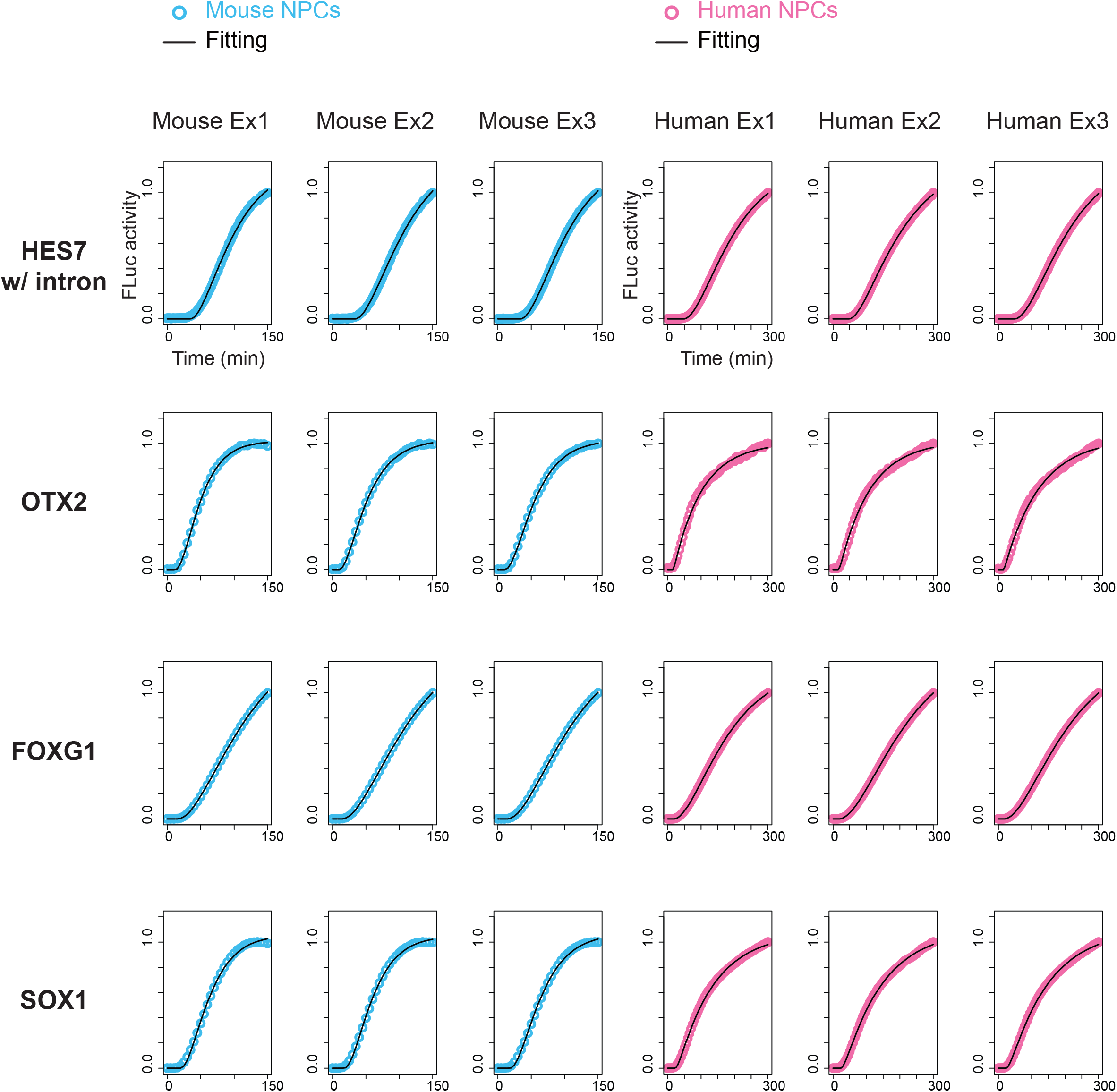
Measurements of the delays in NPCs. Fitting of the delays in NPCs. The original data are shown in Fig. 4j.

## Supplementary Text 1: Parameter dependency of simulated oscillation periods

The mathematical model of the HES7 system that we use to describe the behavior of the segmentation clock is [1]:

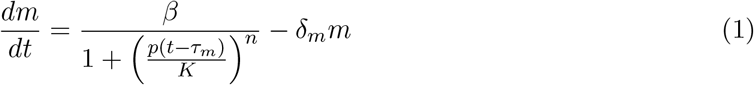

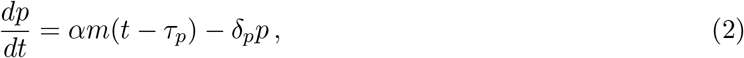

where *m* and *p* are the concentrations of HES7 mRNA and protein, respectively, *α* and *β* are the translation and transcription rates, *K* is the repression threshold, *n* is the repression Hill coefficient, *δ_m_* and *δ_p_* are the degradation rates of the mRNA and protein, which we have measured experimentally as explained in the main text. The mRNA delay *τ_m_* is composed by the repression delay *τ*_Rp_, the transcription delay *τ*_Tx_, and the intron delay *τ*_In_, all of which we have measured experimentally as well. The protein delay *τ_p_*, in turn, corresponds to the translation delay *τ*_Tl_, which was also quantified using our experimental observations.

This system has a fixed point (*m^∗^, p^∗^*) for which the two derivatives above are zero, which obeys:

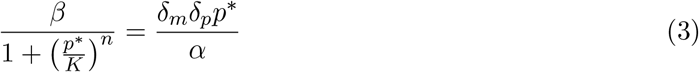

The stability of this fixed point can be analyzed by assuming the following temporal response to a small perturbation (*a, b*):

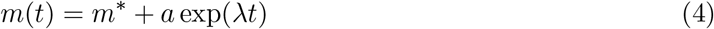

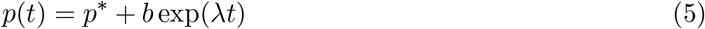

Introducing expressions (4)-(5) into Eqs. (1)–(2), linearizing around *a* = *b* = 0, and imposing that a solution of the form (4)–(5) exists with nonzero *a* and *b*, leads to the following transcendental characteristic equation for the eigenvalues *λ*:

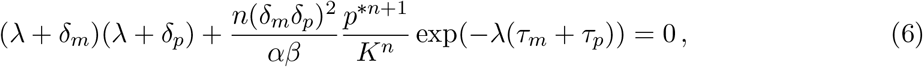

In general the eigenvalues are complex numbers *λ* = *σ* + *iω*. The eigenvalue with highest real part determines the stability of the fixed point, with *σ <* 0 corresponding to an unstable fixed point, and *σ >* 0 to a stable one. The corresponding imaginary part establishes the frequency at which the system oscillates towards the fixed point (if *σ <* 0) or away from it (if *σ >* 0). In the case of an unstable fixed point with *ω* ≠ 0, the system usually falls on a limit cycle whose period can be expected to be close to 2*π/ω*. Taking t hese considerations into account, Eq. (6) s hows t hat **the period of the HES7 oscillations does not depend on the** mRNA **and** protein **delays separately, but only on the total delay** *τ_m_* +*τ_p_*.

For the parameters that we consider in this paper, *p^∗^ ≫ K*, as can be seen in Fig. 1, which represents graphically the solution of Eq. (3) as the crossing point between its left-hand side (blue line) and right-hand side (orange line). In the limit *p^∗^ ≫ K*, Eq. (3) has the following approximate solution:

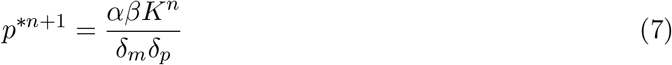

Inserting expression (7) into the characteristic equation (6) leads to:

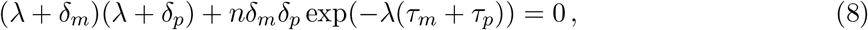

**Figure 1.**
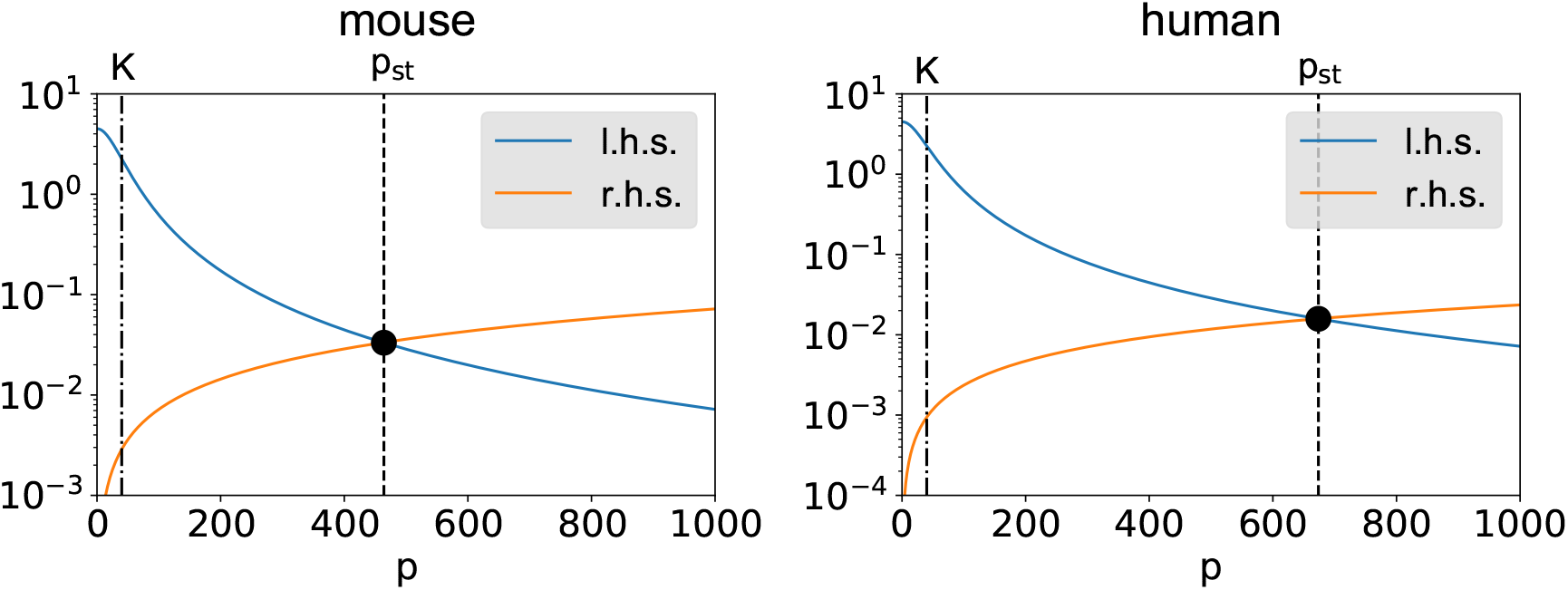
Graphical determination of the fixed point of the HES7 model used in this paper, for the parameters corresponding to the mouse (left) and human (right) cells.

Considering again that the imaginary part of the eigenvalue with the highest real part gives us an estimate of the oscillation period, we can observe from Eq. (8) that **the period of the HES7 oscillations does not depend on the values of the translation and transcription rates** *α* **and** *β***, nor on the repression threshold** *K*.

Finally, if we focus on the bifurcation point (*σ* = 0), we can obtain in closed form the period of the oscillations at that point by computing *ω*. To that end, we write the real and imaginary parts of Eq. (8) for *λ* = *iω* and divide one by the other, to reach the following transcendental equation:

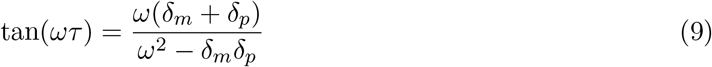

We can thus see that **at the bifurcation point, the period of the oscillations does not depend on** *n***, but only on the degradation rates of the mRNA and the protein, and on the total delay**. These observations are reproduced by our numerical simulations.

**Supplementary Table 1 Biochemical parameters of HES7**

**Supplementary Table 2 Genetic constructs**

**Supplementary Video 1** Time-lapse imaging of HES7 reporter activity in murine (left) and human (right) PSM.

**Supplementary Video 2** Time-lapse imaging of HES7 reporter activity of a single PSM cell in a sparse culture.

